# Control of pili synthesis and putrescine homeostasis in *Escherichia coli*

**DOI:** 10.1101/2024.08.13.607059

**Authors:** Iti Mehta, Jacob Hogins, Sydney Hall, Gabrielle Vragel, Sankalya Ambagaspitiye, Philippe Zimmern, Larry Reitzer

## Abstract

Polyamines are biologically ubiquitous cations that bind to nucleic acids, ribosomes, and phospholipids and, thereby, modulate numerous processes, including surface motility in *Escherichia coli*. We characterized the metabolic pathways that contribute to polyamine-dependent control of surface motility in the commonly used strain W3110 and the transcriptome of a mutant lacking a putrescine synthetic pathway that was required for surface motility. Genetic analysis showed that surface motility required type 1 pili, the simultaneous presence of two independent putrescine anabolic pathways, and modulation by putrescine transport and catabolism. An immunological assay for FimA—the major pili subunit, reverse transcription quantitative PCR of *fimA*, and transmission electron microscopy confirmed that pili synthesis required putrescine. Comparative RNAseq analysis of a wild type and Δ*speB* mutant which exhibits impaired pili synthesis showed that the latter had fewer transcripts for pili structural genes and for *fimB* which codes for the phase variation recombinase that orients the *fim* operon promoter in the ON phase, although loss of *speB* did not affect the promoter orientation. Results from the RNAseq analysis also suggested (a) changes in transcripts for several transcription factor genes that affect *fim* operon expression, (b) compensatory mechanisms for low putrescine which implies a putrescine homeostatic network, and (c) decreased transcripts of genes for oxidative energy metabolism and iron transport which a previous genetic analysis suggests may be sufficient to account for the pili defect in putrescine synthesis mutants. We conclude that pili synthesis requires putrescine and putrescine concentration is controlled by a complex homeostatic network that includes the genes of oxidative energy metabolism.

## Introduction

Polyamines are flexible aliphatic cations that are found in virtually all organisms. *E. coli* contains putrescine (1,4-diaminobutane) and lesser amounts of spermidine (N-(3-aminopropyl)-1,4-diaminobutane), and cadaverine (1,5-diaminopentane) (1, 2). Both the pathways and enzymes of polyamine synthesis are redundant (Figure 1). Strains that lack eight of these genes do not contain detectable polyamines, grow slowly aerobically, and do not grow anaerobically (3). Polyamines interact with nucleic acids and phospholipids, can affect chromosome and ribosome structure, ribosome-mRNA interactions, and protein and nucleic acid elongation rates (4). Polyamines modulate, but are not required for, protein synthesis (5) and expression of hundreds of genes (6). A major mechanism of polyamine-dependent regulation is translational control of genes for 18 major transcription regulators: four of the seven σ factors – σ^18^ (FecI), σ^24^ (RpoE), σ^38^ (RpoS), and σ^54^ (RpoN); three histone-like DNA-binding proteins (Fis, H-NS, and StpA); and CpxR, Cya, Cra, and UvrY (5, 6).

**Figure 1.**
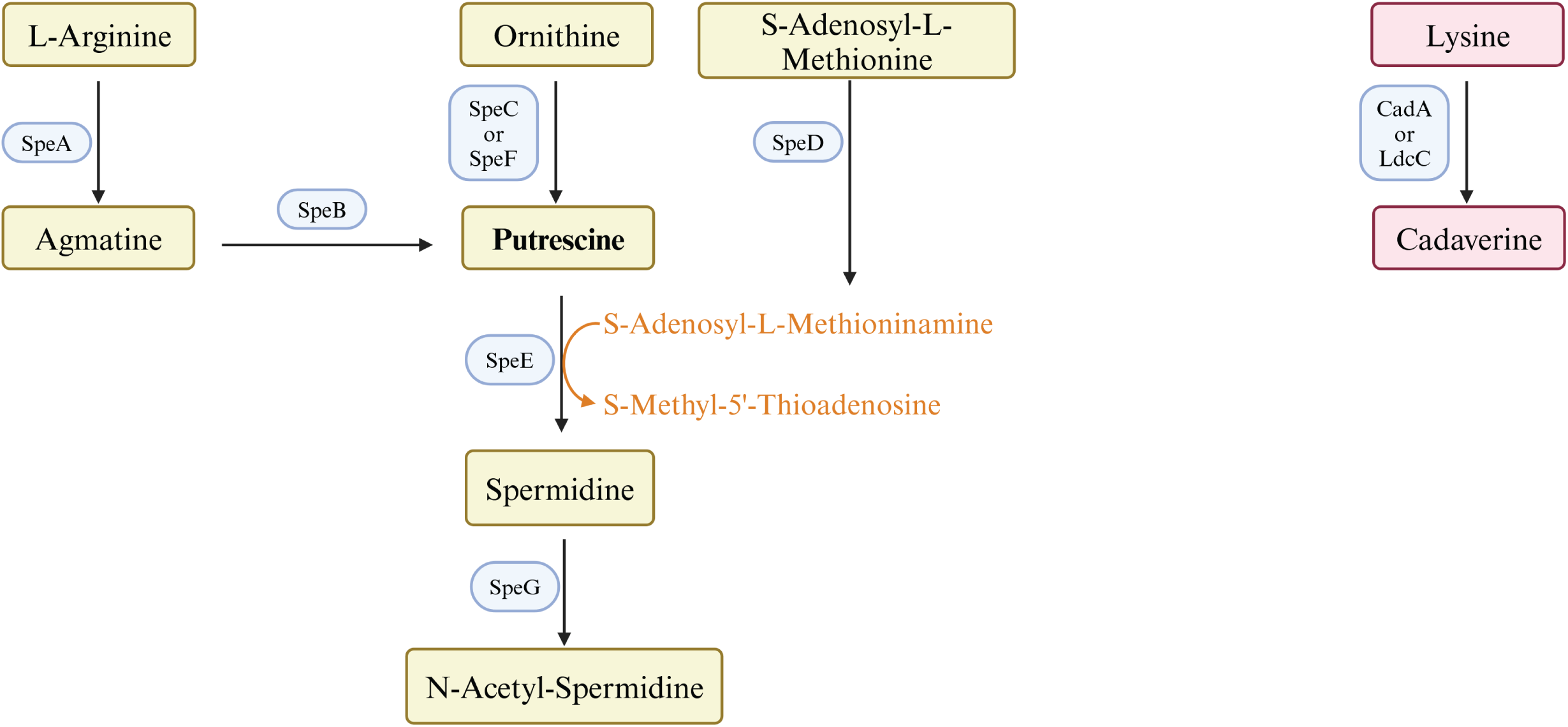
Pathways and enzymes of polyamine synthesis.

Polyamines control flagella-dependent surface motility of *Proteus mirabilis* and pili-dependent surface motility (PDSM) of *E. coli*: the latter has been reported to require the polyamine spermidine (7, 8). While reconstructing putrescine-deficient strains to characterize the basis for the anaerobic growth defect, we noticed that a mutant unable to synthesize spermidine moved normally, while mutants lacking enzymes of putrescine synthesis moved poorly. We characterized the role of polyamines on *E. coli* surface motility and pili synthesis by genetic analysis, ELISA assays, RT-qPCR, electron microscopy, and RNAseq. We showed that PDSM and pili synthesis required two independent pathways of putrescine synthesis, but did not require spermidine, and was modulated by putrescine transport and catabolism. Results from an RNAseq analysis are consistent with multiple mechanisms of putrescine-dependent control, and, unexpectedly, suggests a putrescine homeostatic network that rewires metabolism to maintain intracellular putrescine.

## Results

### W3110 surface motility requires pili

An assessment of surface motility for *E. coli* K-12 and eight derivatives showed five strains covered a plate in 12-18 hours, while four did not traverse a plate in 36 hours (9). The slow-moving strains generated genetically stable fast-moving variants which suggests that the former were ancestral. We chose our lab strain of W3110 for further study because (a) it exhibited the least variability in spreading diameter and generated fewer fast-moving variants, and (b) we had previously analyzed this strain for the energy requirements for surface motility (10). We refer to our slow-moving lab strain hereafter as W3110 but not all W3110 strains move similarly: the W3110 from the genetic stock center is a fast-moving variant (9).

Several results suggested that W3110 requires pili for surface motility. First, deletion of *fimA*, which codes for the major type 1 pili subunit, abolished the oscillatory pattern and reduced, but did not eliminate, surface motility, but deletion of *fliC* which encodes the major flagellum subunit did not affect surface motility (Figure 2A). Second, electron microscopy confirmed that W3110 isolated from surface motility plates possessed pili (Figure 2B). None of >500 cells expressed flagella. A mixed population of elongated (3-4 µm) and non-elongated (<2 µm) cells were visible, and pili were mostly associated with elongated cells. Finally, W3110 surface motility requires glucose (10) which should prevent flagella synthesis via inhibition of cyclic AMP synthesis (11). We confirmed that glucose suppressed swimming motility which implies suppression of flagella synthesis in W3110 during surface motility assays (Figure 2C). A W3110 derivative with deletions of both *fliC* and *fimA* still exhibited outward movement on a surface motility plate (Figure 2A), but electron microscopy showed the absence of appendages (Figure 2C). In summary, genetics, electron microscopy, and a glucose requirement show that W3110 surface motility requires pili.

**Figure 2.**
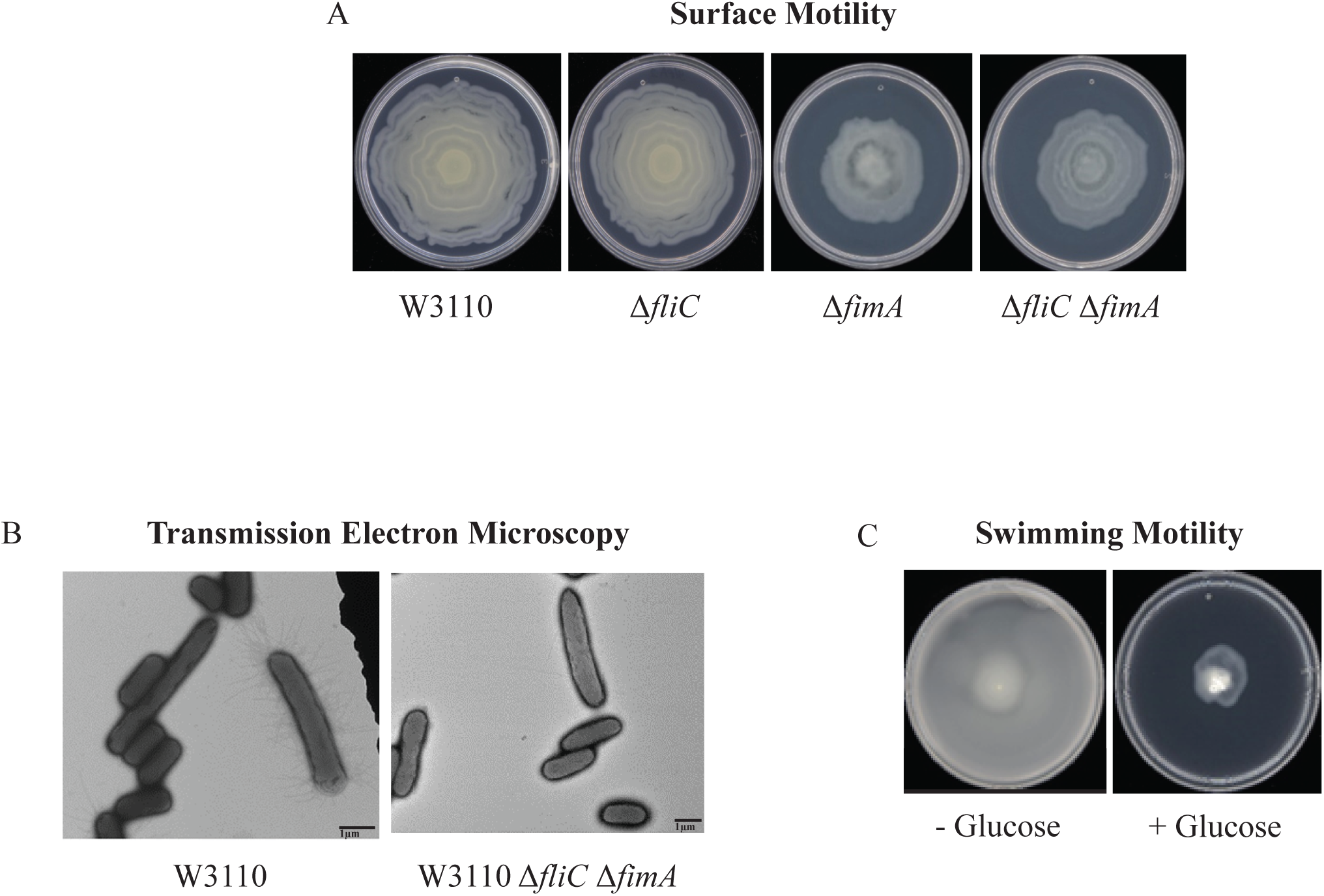
Surface motility of W3110. (A) Surface motility of parental and derivative strains lacking the major subunits of the flagella (FliC), pili (FimA), or both. (B) TEM images of cells of W3110 and W3110 Δ*fliC* Δ*fimA* taken from the movement’s edge directly from surface motility plates. (C) Swimming motility with and without 0.5% glucose.

Results from W3110 sequencing found an *IS1* insertion in *fimE* which could explain its greater movement that is less variable and greater genetic stability. The FimB and FimE recombinases are the major components of phase variation that determines the orientation of the promoter that initiates transcription of the *fimAICDFGH* operon which codes for the pili structural proteins. The FimB recombinase favors the productive ON orientation, while the more active FimE recombinase favors the unproductive OFF orientation (12–14). The insertion in *fimE* is the likely basis for the relative stability of W3110.

### PDSM required two independent putrescine synthesis pathways

We examined PDSM requirements for polyamine anabolic pathways. A major putrescine biosynthetic pathway is ornithine decarboxylation either by the constitutive SpeC or the low-pH inducible SpeF. The Δ*speC* and Δ*speF* mutants moved as well as the parental strain, but a Δ*speC* Δ*speF* double mutant moved less well (Figure 3). SpeA (arginine decarboxylase) and SpeB (agmatinase) catalyze an alternate two-step putrescine synthetic pathway (Figure 1) (15): deletion of either gene reduced surface motility (Figure 3). The D*speA*, D*speB*, and D*speC* D*speF* strains grew as well as the parental strain (Supplemental Figure 1A) and had normal swimming motility (Supplemental Figure 1B). A *ΔspeE* mutant, which lacks the enzyme for spermidine synthesis, moved on a surface as well as the parental strain (Figure 3).

**Figure 3.**
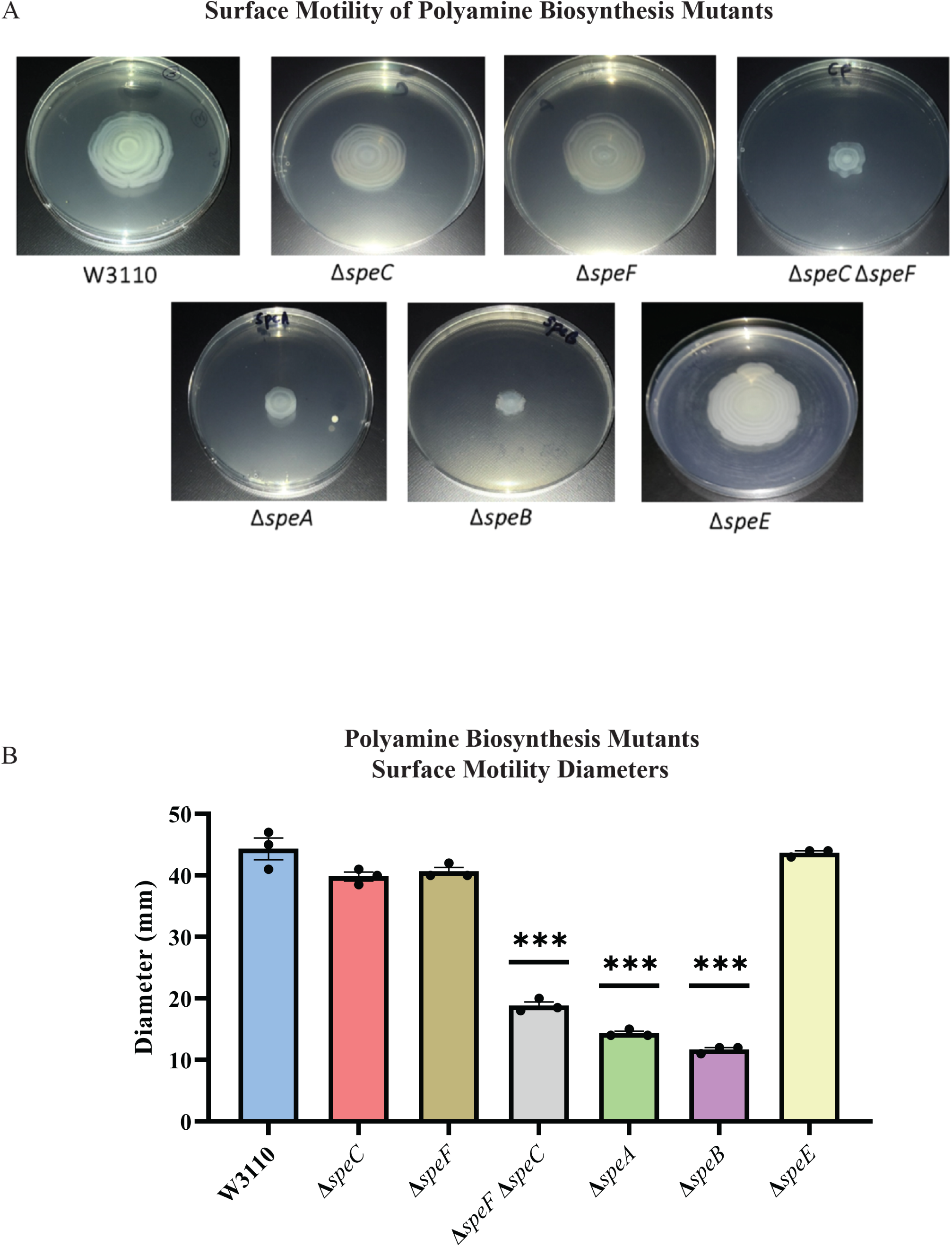
Genetics of surface motility. (A) PDSM of mutants with defects in polyamine anabolic genes. All assays were performed in triplicate, and representative images are shown. (B) Diameter of surface movement of polyamine mutants after 36 hours. Error bars represent standard deviations for three independent replicates. Statistical analysis was performed using the Dunnett test of significance: *, *P*<0.05; **, *P*<0.01; ***, *P*<0.001. In this figure the Δ*speB* mutant was IM26.

Supplemental putrescine at 1 mM restored surface motility of a *speB* mutant (Figure 4A) and a *speA* mutant (not shown). Spermidine did not restore wild type movement and, also, caused an unusual pattern of movement in the parental strain (Figure 4A). These exogenous concentrations of putrescine and spermidine had no effect on growth rates in liquid motility medium (not shown). In summary, PDSM required the simultaneous presence of two independent pathways of putrescine synthesis but did not require spermidine.

**Figure 4:**
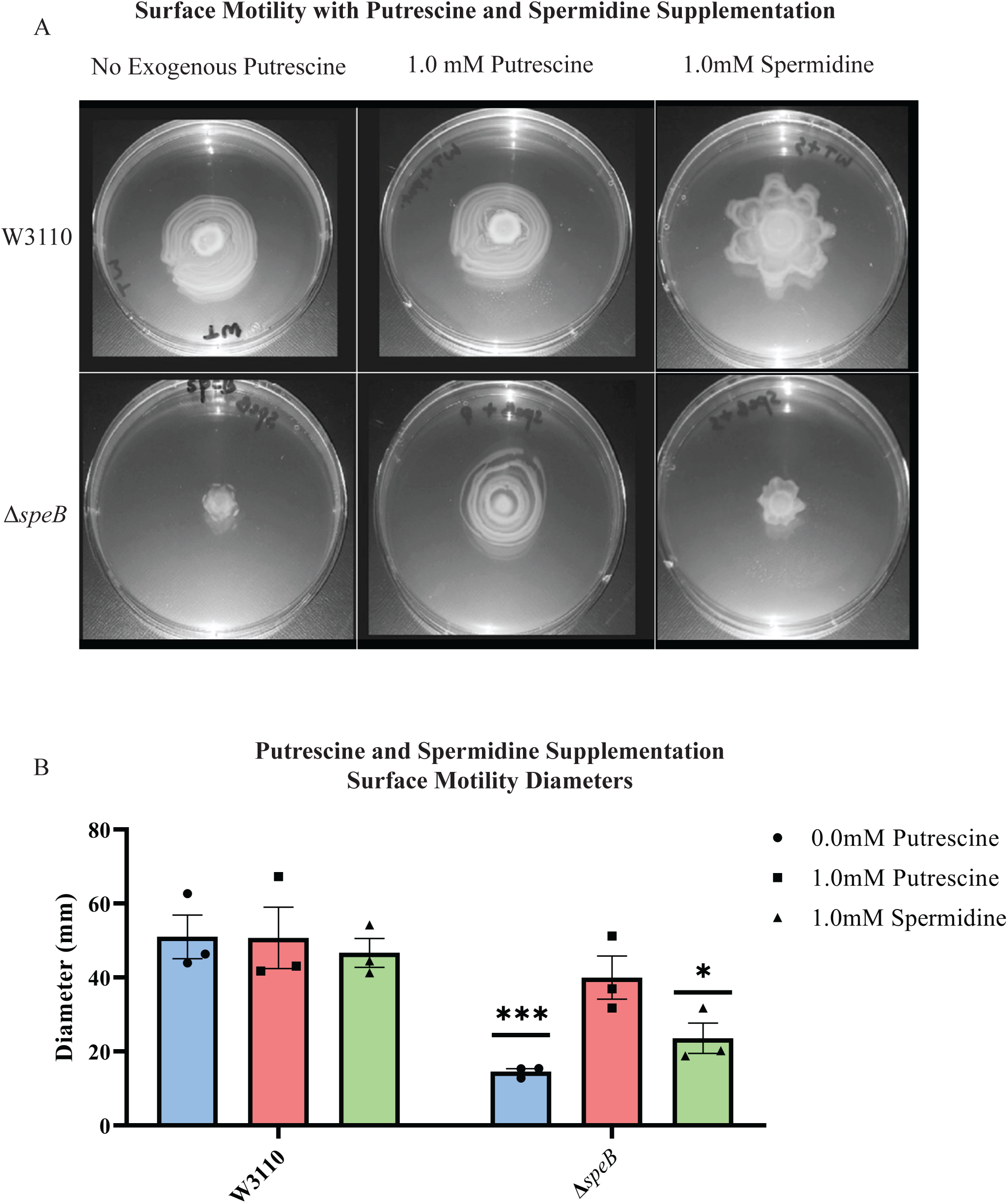
Nutritional supplementation of Δ*speB* mutant. (A) Putrescine and spermidine supplementation of wild type and Δ*speB* strains. (B) Diameter of surface movement of polyamine mutants after 36 hours. Error bars represent standard deviations for three independent replicates. Statistical analysis was performed using the Sadik test of significance: *, *P*<0.05; **, *P*<0.01; ***, *P*<0.001. All assays were performed in triplicates and representative images are shown. In this figure the Δ*speB* mutant was IM26.

### Loss of putrescine transport systems affected surface motility

*E. coli* has several polyamine transport systems and one has been reported to require for surface motility (16, 17). Mutants lacking the PlaP and PotF putrescine transporters had 50% and 40% reduced surface motility, respectively, and lost the concentric ring pattern (Figure 5A). Loss of the PotE and PuuP transporters affected neither the movement diameter (Figure 5B) nor concentric ring formation (Figure 5A). We note that *potF* and *plaP* are more highly expressed than *potE* and *puuP* (85, 64, 17, and 6 counts per million transcripts, respectively) for W3110 grown in liquid motility medium (18). We conclude that extracellular putrescine and its transport contribute to PDSM.

**Figure 5:**
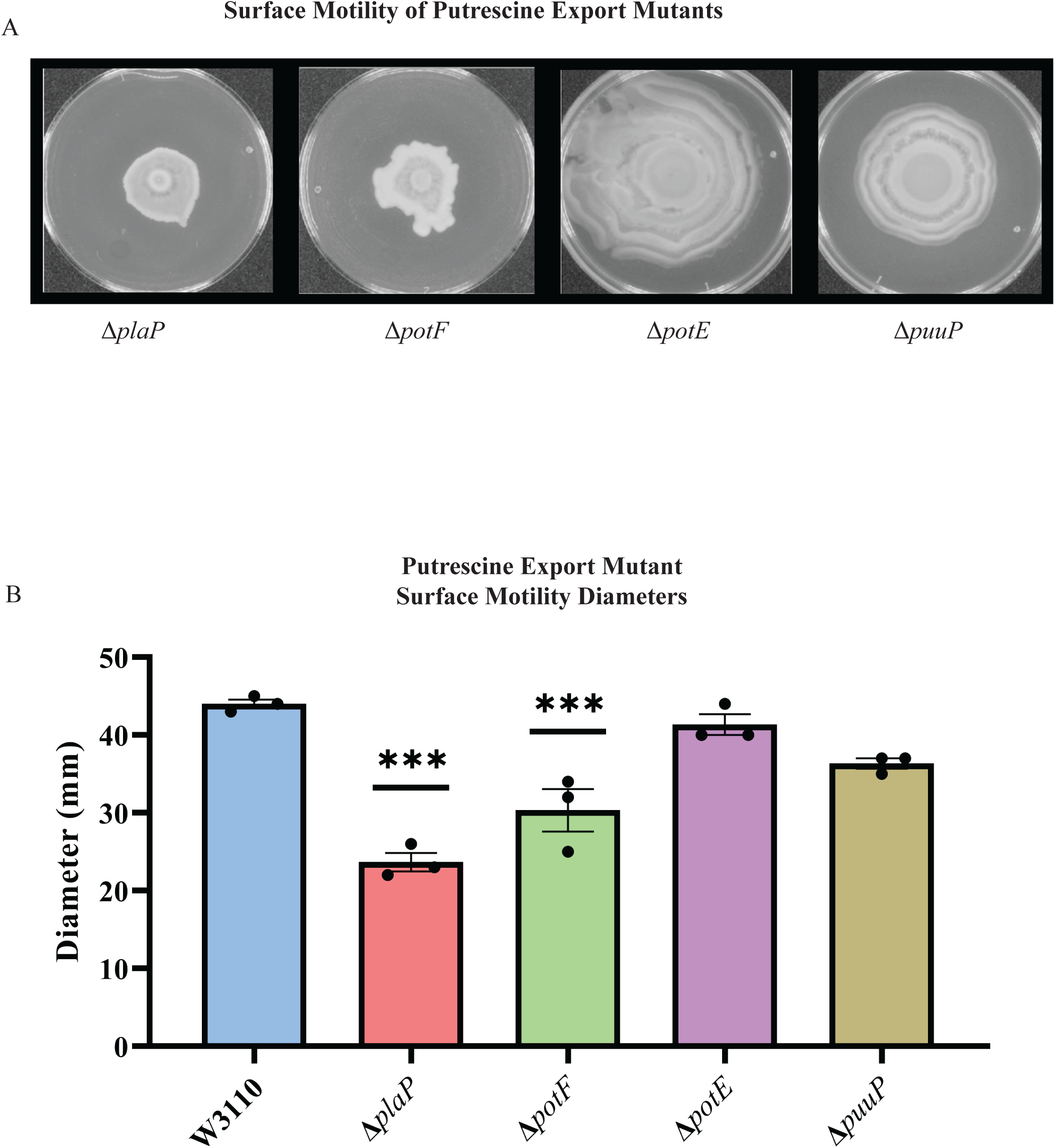
Putrescine transport and surface motility. (A) Representative images of surface motility for strains defective in putrescine transport genes. (B) Diameter of surface movement of polyamine mutants after 36 hours. Error bars represent standard deviations for three independent replicates. Statistical analysis was performed using the Dunnett test of significance: *, *P*<0.05; **, *P*<0.01; ***, *P*<0.001. All assays were performed in triplicate and representative images are shown.

### Putrescine catabolism affected surface motility

Because putrescine catabolism modulates intracellular putrescine concentrations (19, 20), we examined PDSM in putrescine catabolic mutants. The major putrescine catabolic pathway is initiated with putrescine glutamylation (Figure 6A, right column), and a second pathway initiates with putrescine deamination (Figure 6A, left column) (20). Deletion of either *puuA* or *patA* which code for the first enzymes of their respective pathways did not affect surface motility (Figure 6B). A double mutant moved less well which was unexpected because the proposed higher intracellular putrescine should have stimulated PDSM. Consistent with the stimulatory effect of putrescine, loss of *puuR* which codes for the repressor of PuuA-initiated pathway genes (20), impaired PDSM (Figure 6B). One possible explanation to reconcile these seemingly contradictory results is that an optimal polyamine concentration stimulates PDSM, and a high concentration is inhibitory. The next section describes experiments to test this possibility.

**Figure 6:**
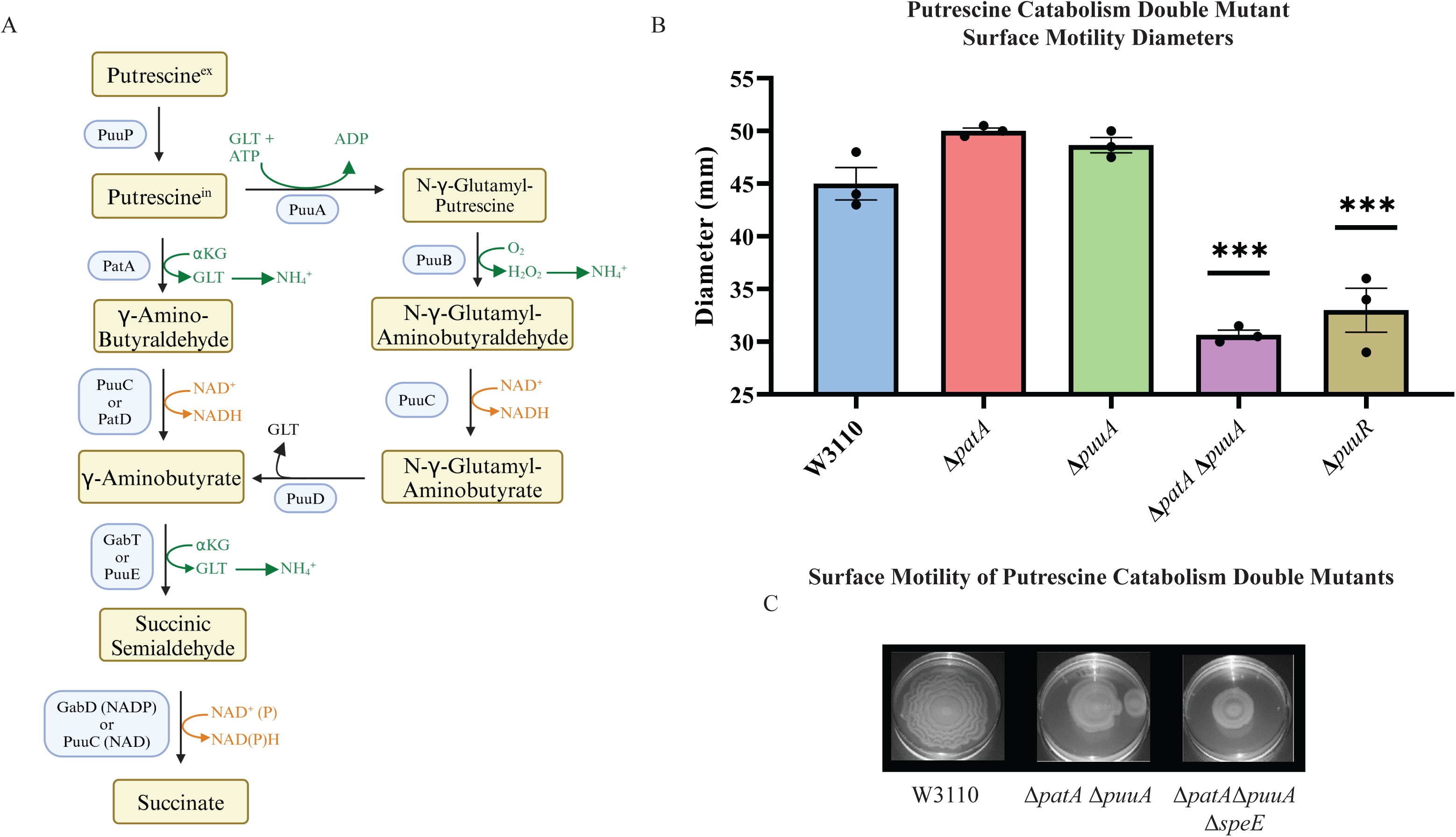
Putrescine catabolism and surface motility. (A) Pathways and enzymes of putrescine catabolism. (B) Motility diameter of putrescine catabolic mutants after 36 hours. Error bars represent standard deviations for three independent replicates. Statistical analysis was performed using the Dunnett test of significance: *, *P*<0.05; **, *P*<0.01; ***, *P*<0.001. All assays were performed in triplicates and representative images are shown. (C) Effect of loss of *speE* on the *patA puuA* catabolic double mutant.

### High putrescine reduced expression of pili genes

High putrescine inhibits translation of some mRNAs, and high spermidine, a product of putrescine metabolism, inhibits growth (21, 22). If the phenotype of the Δ*patA* Δ*puuA* double mutant results from spermidine toxicity, then loss of spermidine synthase (SpeE) should reverse the phenotype. This prediction was not met which argues against spermidine toxicity (Figure 6C). For the *speB* mutant, 1 mM putrescine stimulated PDSM, and 4 mM putrescine was slightly inhibitory (Figure 7A and B). The concentric ring pattern was observed with 1 mM, but not 4 mM putrescine. RT-qPCR to test transcriptional regulation of pili expression by putrescine showed optimal *fimA* transcription at 1 mM putrescine in the *speB* mutant (Figure 7C). Indirect enzyme linked immunosorbent assays (ELISAs) against FimA suggested that the level of FimA in the *speB* mutant at 1 mM putrescine was higher than with 4 mM putrescine, but the difference was not statistically significant (Figure 7D). Variation of supplemental putrescine did not affect FimA in the parental strain (Figure 7C and 7D). Transmission electron microscopy (TEM) showed that pili expression was highest for the *speB* mutant at 1 mM putrescine (Figure 8), moderate at 0.1- and 4-mM putrescine, and undetectable without putrescine. The cells grown with 4 mM putrescine were shorter and thinner, which suggests stress, possibly mimicking osmotic stress. From these results and those in the previous section, we conclude that pili expression required an optimal putrescine concentration.

**Figure 7:**
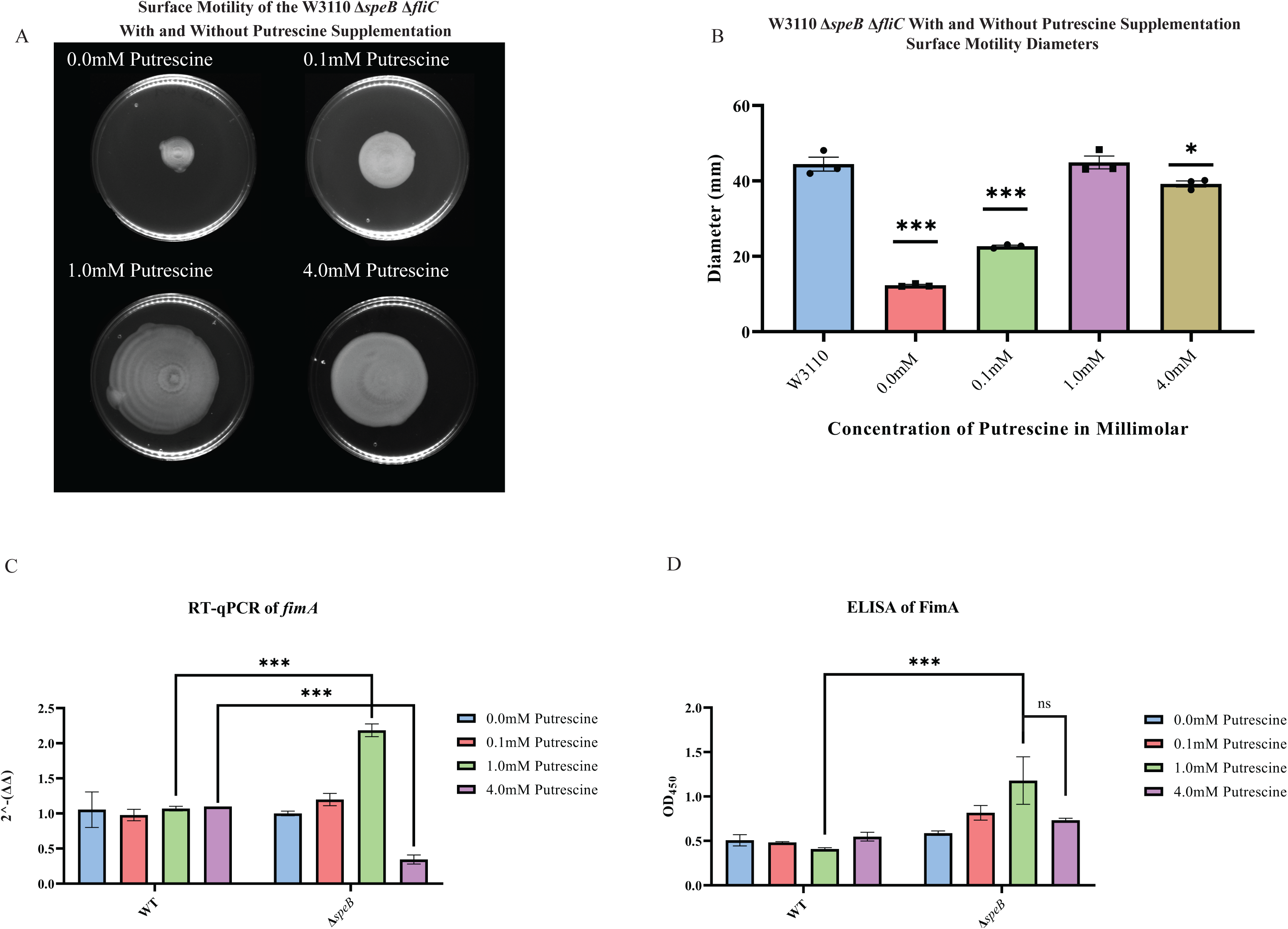
Surface motility and pili expression in W3110 and Δ*speB*. (A) W3110 Δ*speB* surface motility with 0.0, 0.1, 1.0, and 4.0 mM exogenous putrescine. All assays were performed in triplicate. Representative images are shown. (B) Average diameter of 3 separate surface motility plates. The parental strain without putrescine is shown for reference. Significance was determined by comparing the diameters of the Δ*speB* mutants in the different concentrations of putrescine compared to the parental W3110. One way ANOVA was used with Dunnett hypothesis testing to determine p values, ***, *p <0.001.* (C) Reverse transcriptase-quantitative PCR using primers targeting the *fimA* gene from cells grown in 0.0, 0.1, 1.0, and 4.0 mM of supplemented putrescine. Double deltas were generated by normalizing the parental and the Δ*speB* mutant RNA libraries to *rpoD* then comparing *fimA* expression. Two-way ANOVA was used to determine significance followed by Sadik hypothesis testing. ***, *p*<0.001. (D) Enzyme linked immunosorbent assays using antibodies targeting pili (FimA) from cells grown with 0.0, 0.1, 1.0, and 4.0 mM of supplemented putrescine. Two-way ANOVA was used to determine significance followed by FDR adjusting. ***, *p*<0.001. In this figure the Δ*speB* mutant was J15.

**Figure 8:**
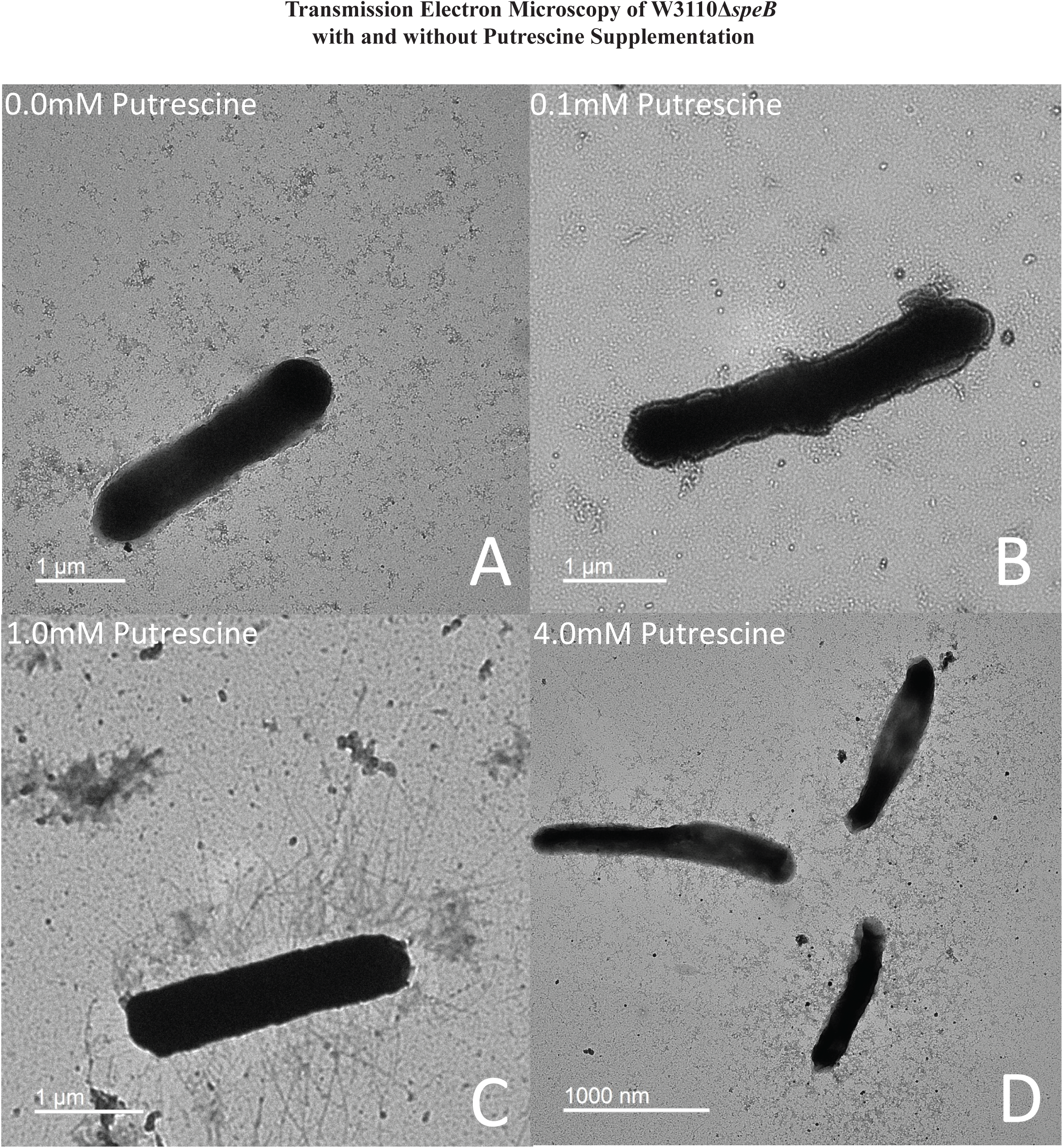
Transmission electron micrographs of Δ*speB* mutant cells after surface motility. Representative images are shown and cells were removed from motility plates with the following exogenous putrescine concentrations: (A) none, (B) 0.1 mM, (C) 1.0 mM, and (D) 4.0 mM. No pili were observed with 0 and 0.1 mM putrescine. Optimal pili production was observed with 1.0 mM putrescine. The bar represents one micron. In this figure the Δ*speB* mutant was J15.

### RNAseq analysis identifies *fim* operon expression and energy metabolism as targets of putrescine control

RNAseq analysis was used to further confirm or determine whether putrescine affected (a) transcription of the *fim* operon that codes for the pili structural genes, (b) *fim* operon promoter orientation, i.e. phase variation, or (c) energy metabolism: surface motility of W3110 has been shown to require glucose metabolism and oxidative phosphorylation (10).

#### RNAseq analysis

We grew parental W3110 and its Δ*speB* derivative with and without 1 mM putrescine. *R^2^* values of pairwise comparisons showed that the transcriptome of the *speB* mutant grown without putrescine differed from the other transcriptomes which were similar (Table 1 and Supplemental Figure 2). A multidimensional scaling plot and a plot of the 100 most variable genes visualize these comparisons (Figure 9A and B). When compared to the parental strain, the *speB* mutant grown without putrescine had 310 downregulated genes and 159 upregulated genes with at least four-fold differential expression and FDR < 0.05. Transcripts for the putrescine-induced *puuAP* and *puuDRCBE* operons, which specify genes of the major putrescine catabolic pathway, were reduced from 1.6- to 14-fold fold (FDR ≤ 0.02) in the *speB* mutant (Supplemental Table 1). Because higher intracellular putrescine results in higher catabolic gene expression, these results imply lower intracellular putrescine in the Δ*speB* mutant.

**Figure 9:**
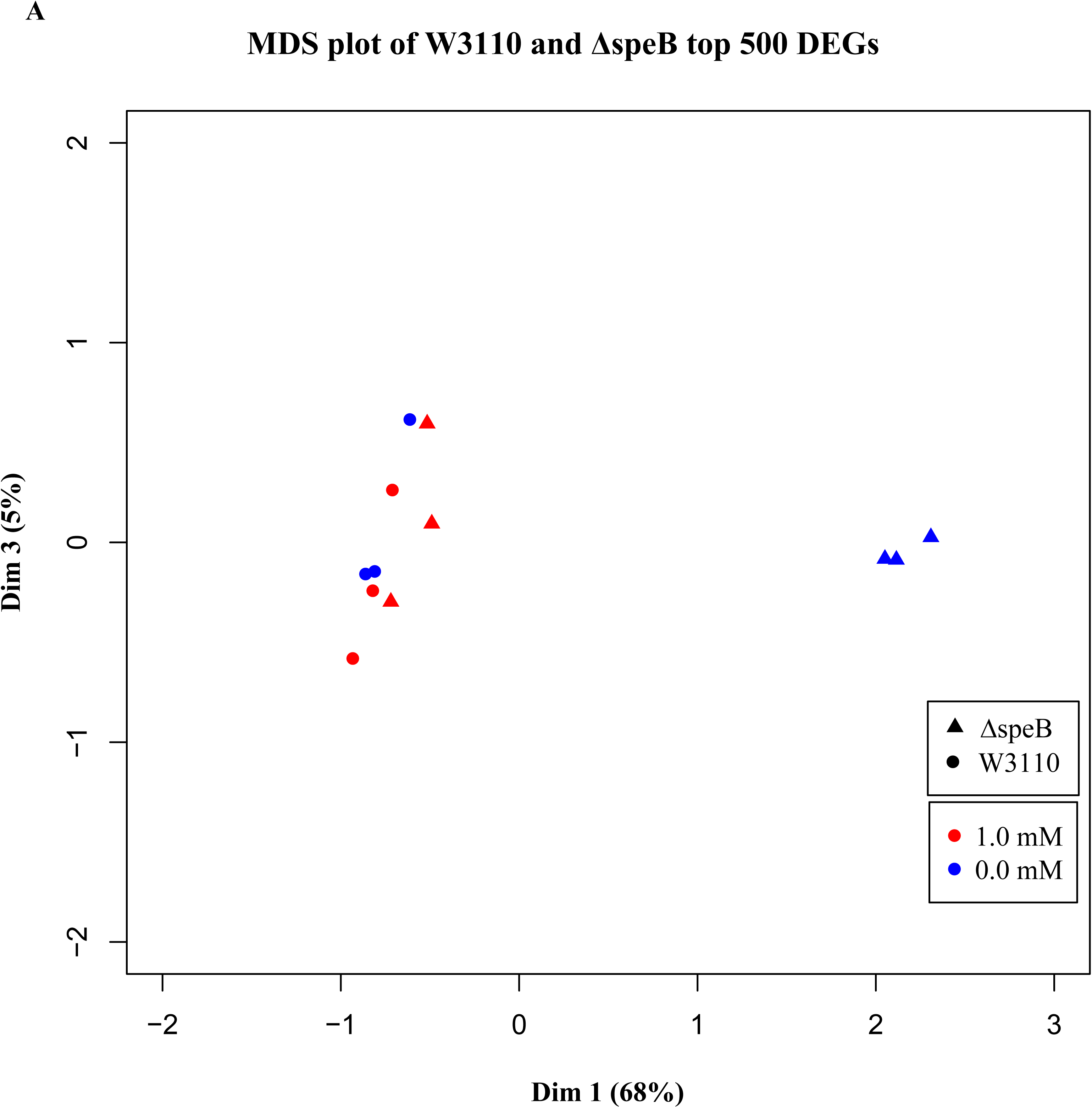
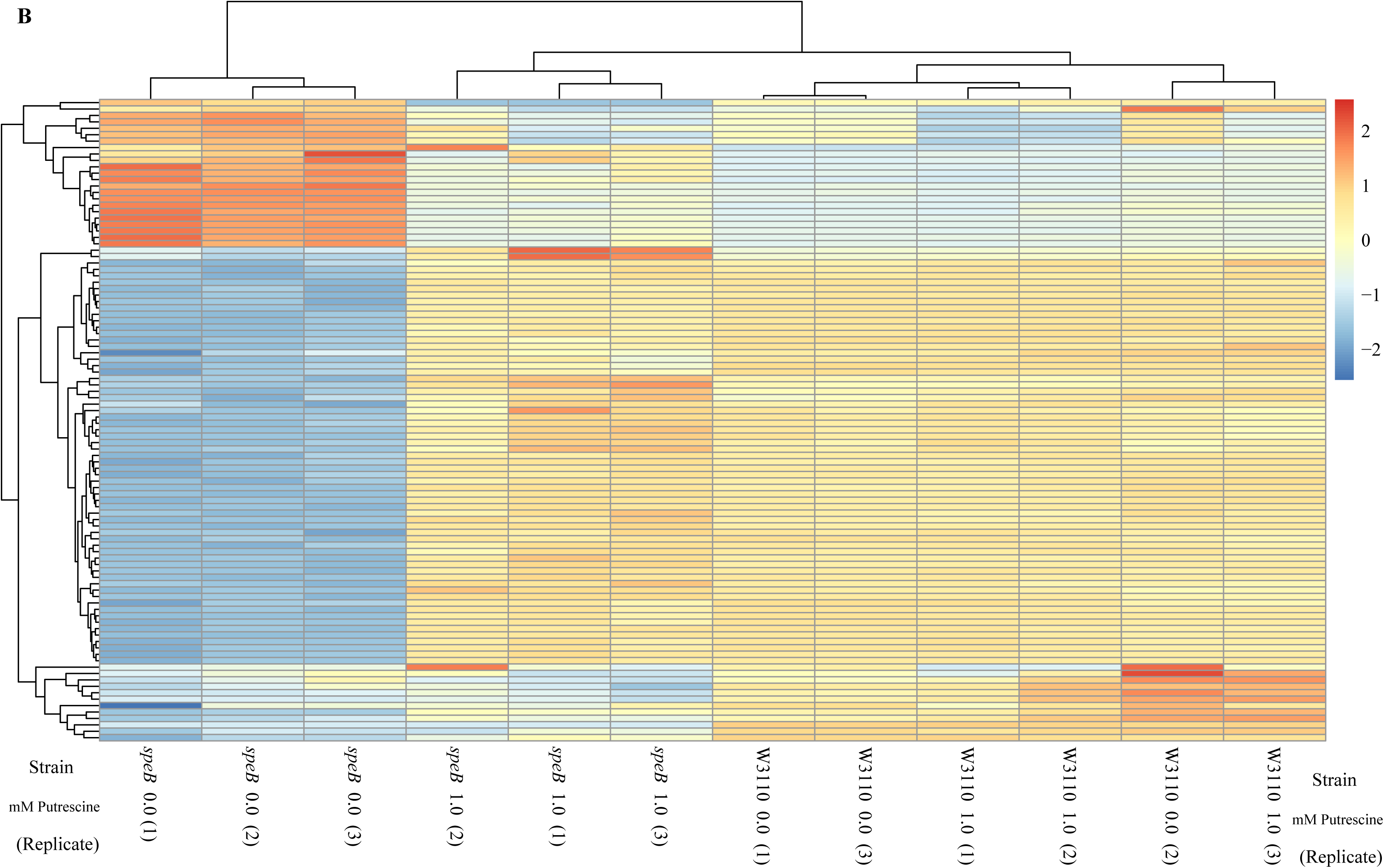
Visual representation of the results from transcriptomic sequencing of W3110 and the Δ*speB* mutant’s gene expression in media with and without 1.0 mM putrescine. (A) Multidimensional scaling plot of the parental and Δ*speB* mutant transcriptomes. When grown with 1.0 mM putrescine (red), the Δ*speB* mutant transcriptome (triangles) are nearly identical to the parental transcriptomes when grown without and with putrescine (blue and red circles, respectively). When grown without putrescine, the Δ*speB* mutant transcriptome (blue triangles) is greatly skewed from transcriptomes of the parental strain and the Δ*speB* mutant grown with putrescine. (B) Heatmap of the top 100 most variable genes further demonstrates the distinctiveness of the Δ*speB* mutant grown without putrescine and the similarities of the Δ*speB* mutant transcriptome when grown with 1.0 mM putrescine supplementation and the parental grown with or without putrescine. In this figure the Δ*speB* mutant was J15.

**Table 1.** Pairwise statistical comparisons of transcriptomes. Abbreviation: putr is putrescine.

| Strain 1 | Strain 2 | $R^2$ |
| --- | --- | --- |
| Wild-type (1 mM putr) | wild-type (0 mM putr) | 0.950 |
| Wild-type (1 mM putr) | $\Delta speB$ (1 mM putr) | 0.954 |
| $\Delta speB$ (0 mM putr) | $\Delta speB$ (1 mM putr) | 0.820 |
| $\Delta speB$ (0 mM putr) | wild-type (0 mM putr) | 0.796 |
| $\Delta speB$ (0 mM putr) | wild-type (1 mM putr) | 0.786 |

#### Transcription of the *fim* operon is reduced in the Δ*speB* mutant

Deletion of *speB* reduced transcripts for genes of the *fimA* operon (Figure 10A). Numerous transcription factors activate the *fim* operon (23, 24), and, of these, loss of *speB* reduced *hns* transcripts, increased *fis*, *lrp*, and *qseB* transcripts, but had no effect on *ihfA* and *ihfB* (Figure 10B). Loss of *hns*, *ihfA*, and *lrp* impaired PDSM (Figure 10C and D), which confirms the requirement for their products for pili synthesis in our strain background.

**Figure 10:**
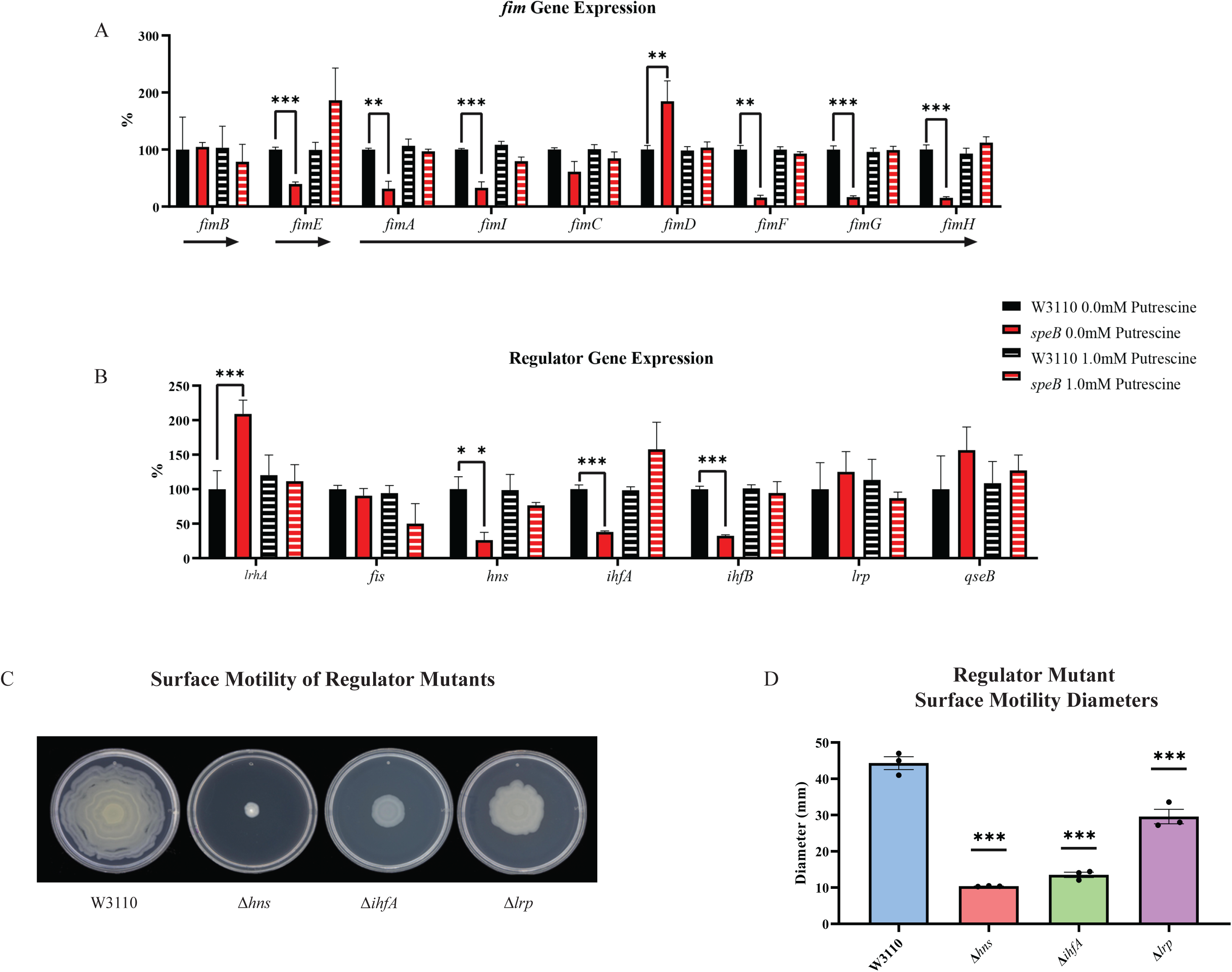
Transcriptomic sequencing of genes for the *fim* region and regulators that control *fim* gene expression. (A) Expression of the *fim* genes in the parental W3110 and the *speB* mutant with and without putrescine supplementation. Values were calculated by dividing the individual replicates’ counts per million (CPM) value for each gene by the mean CPM of that gene in W3110 grown without putrescine and multiplying by 100 to yield the percent expression. Arrows below the genes signify the known operons: *fimB* and *fimE* belong to single gene operons, while *fimAICDFGH* belong to one operon. One way ANOVA was used to determine significance using Dunnett hypothesis testing. **, *p* <0.01; ***, *p* <0.001. (B) Expression of some regulators known to affect *fim* gene expression. Values were calculated as described in (A). ***, *p* <0.001. (C) Surface motility of W3110 and three regulator mutants (Δ*hns*, Δ*ihfA*, and Δ*lrp*). A gene found to be significantly different by this transcriptomic analysis (*hns*) was confirmed to be important in surface motility. In this figure the Δ*speB* mutant was J15. (D) Diameter of surface movement of regulatory mutants after 36 hours. Error bars represent standard deviations for three independent replicates. Statistical analysis was performed using the Dunnett test of significance: *, *P*<0.05; **, *P*<0.01; ***, *P*<0.001. All assays were performed in triplicate.

#### Putrescine does not control *fim* operon phase variation in W3110. The phase ON-favoring

FimB and phase OFF-favoring FimE recombinases control the orientation of the *fim* operon promoter. Our version of W3110 has an insertion in *fimE* which means that FimB is the major factor that controls phase variation. Loss of *speB* reduced transcripts from *fimB* which could account for loss of motility. Using specific primers, PCR analysis showed that loss of *speB*, loss of *hns*, or the presence of putrescine did not alter the ratio of ON to OFF (Supplemental Figure 3). This experiment determines the steady state phase ON to OFF ratio, which less FimB might not affect if FimB is in excess. In strains with FimE, the rate of phase ON orientation will be much slower and even small changes in FimB activity could conceivably affect the switch orientation. We conclude that putrescine does not control phase variation, at least in our strain of W3110.

#### Evidence for a putrescine homeostatic network

Altered transcript levels in the *speB* mutant suggest compensatory mechanisms for low putrescine (Supplemental Table 1 for transcript differences for all genes, Table 2 for selected genes, and Figure 11 for a diagrammatic representation of transcript differences). The *speB* mutant had more transcripts from (a) genes of arginine and ornithine carboxylase, (b) all genes of ornithine and arginine synthesis from glutamate, and (c) genes for three separate putrescine transport systems; and fewer transcripts from the *sap* operon which codes for a putrescine exporter (25) and *rpmH* and *rpsT*, whose products inhibit ornithine and arginine decarboxylase (26). The net effect of these changes should be to increase intracellular putrescine (solid arrows in Figure 11). In other words, these changes are compensatory for low putrescine. In addition to these changes, the *speB* mutant had 60-fold and 7-fold more transcripts for *mgtA* and *phoBR* which code for the major magnesium transporter and the regulators of phosphate assimilation, respectively. These changes may also be compensatory and are discussed below.

**Figure 11:**
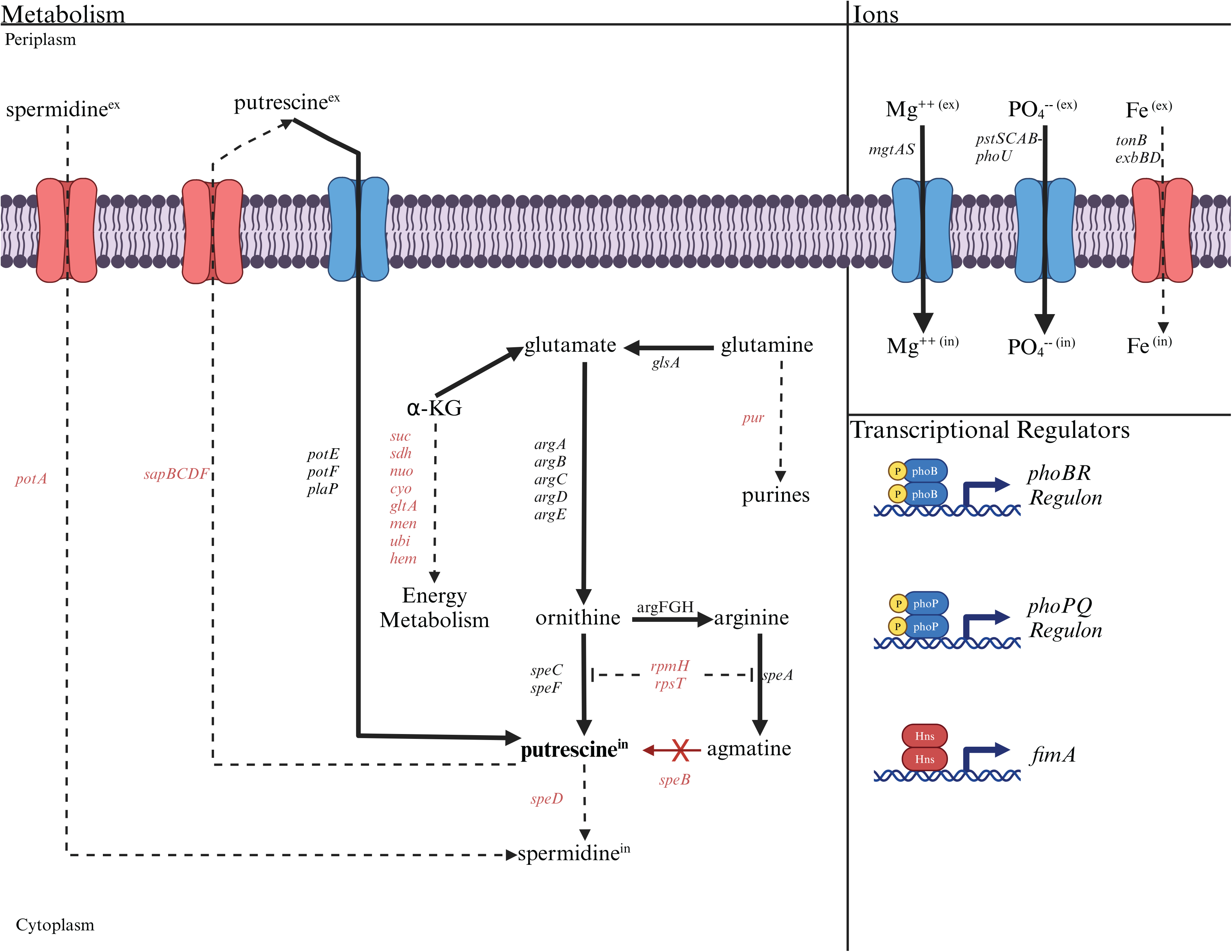
The deduced diversion of metabolism in a *speB* mutant away from energy metabolism toward putrescine synthesis compared to the parental strain. The effects of low putrescine are shown. Genes and processes (transport and transcriptional regulators) in red have fewer transcripts, while those in black or blue have more. The dashed lines represent proposed reduction in metabolic flux because of fewer transcripts from genes coding for the enzymes involved. Operons are shown except for the larger operons or regulons. For example, *pur* is meant to represent the unlinked genes that code for enzymes of purine synthesis. Table 2 or Supplemental Table 1 should be consulted for specific genes and the quantitative change in transcripts.

**Table 2.** Differentially expressed genes that are proposed to contribute to putrescine homeostasis. A positive number means more transcripts in the Δ*speB* (lower putrescine) strain.

| Gene (function) | log<sub>2</sub>FC | FDR |
| --- | --- | --- |
| **Polyamine synthesis** |  |  |
| *speA* (putrescine) | 0.99 | 0.003 |
| *speC* (putrescine) | 1.43 | 3E-4 |
| *speD* (spermidine) | -0.83 | 0.01 |
| *speF* (putrescine) | 1.05 | 0.007 |
| *rpmH* (inhibitor of SpeA and SpeC) | -2.77 | 2E-4 |
| *rpsT* (inhibitor of SpeA and SpeC) | -1.92 | 8E-4 |
| **Polyamine transport** |  |  |
| *potABCD\** (spermidine) | -2.17 | 3E-4 |
| *potE* (putrescine) | 2.07 | 2E-4 |
| *potFGHI\** (putrescine) | 1.41 | 4E-4 |
| *plaP* (putrescine) | 1.11 | 0.005 |
| *sapBCDF\** (putrescine export) | -1.18 | 8E-4 |
| **Arginine synthesis** |  |  |
| *argA_2* (synthesis) | 1.77 | 0.001 |
| *argB* (synthesis) | 1.87 | 0.002 |
| *argC* (synthesis) | 2.86 | 0.005 |
| *argD* (synthesis) | 1.41 | 0.001 |
| *argF* (synthesis) | 1.63 | 0.01 |
| *argG* (synthesis) | 2.32 | 0.002 |
| *argH* (synthesis) | 0.97 | 0.001 |
| *argI* (synthesis) | 1.95 | 0.018 |
| *argR* (repressor of arginine regulon) | -1.35 | 4E-4 |
| **Glutamate generation** |  |  |
| *glsA* (glutamine degradation) | 2.16 | 5E-4 |
| *glnG* (regulation of ammonia assimilation) | -1.59 | 0.022 |
| **TCA cycle/electron transport** |  |  |
| *acnA* (TCA cycle) | -1.04 | 8E-4 |
| *acnB* (TCA cycle) | -1.96 | 2E-4 |
| *fumA* (TCA cycle) | -1.50 | 6E-4 |
| *fumB* (TCA cycle) | -1.49 | 0.003 |
| *gltA* (TCA cycle) | -1.64 | 2E-4 |
| *icdA* (TCA cycle) | -1.32 | 0.002 |
| *sdhCDAB-sucABCD\** (TCA cycle) | -3.88 | 5E-5 |
| *nuo* operon* (electron transport) | -0.82 | 0.004 |
| *menFDHBCE\** (electron transport) | -0.8 | 0.005 |
| *ubiEJB\** (electron transport) | -0.93 | 0.004 |
| *cyoABCD\** (electron transport) | -1.41 | 9E-4 |
| *hemCD\** (heme) | -2.05 | 2E-4 |
| **Iron transport** |  |  |
| *exbBD\** (transport of all iron chelates) | -2.33 | 4E-5 |
| *tonB* (transport of all iron chelates) | -1.9 | 4E-5 |
| *entCEBA\** (enterochelin iron) | -5.2 | 7E-6 |
| *fecABCDE \** (ferric citrate) | -4.4 | 3E-5 |
| *fecIR* (regulators, ferric citrate) | -4.4 | 6E-6 |
| *feoABC\** (ferrous iron) | -3.4 | 7E-4 |
| *fur* (regulator, iron assimilation) | -0.40 | 0.11 |
| **Magnesium and phosphate transport** |  |  |
| *mgtA* (magnesium) | 5.90 | 5E-6 |
| *phoQ* (regulator, magnesium) | 0.79 | 0.009 |
| *pstSCAB\** (phosphate) | 3.61 | 0.01 |
| *phoBR* * (regulator, phosphate assimilation) | 2.80 | 0.02 |

\* Values for transcripts of the first gene of operon are given. Results for other genes of the operon are in Supplemental Table 1

#### Putrescine affects energy metabolism which could account for loss of motility

Loss of *speB* affected transcripts for genes of the major energy metabolism pathways (Table 2): fewer transcripts for genes coding for all citric acid cycle enzymes, NADH dehydrogenase I (the entire *nuo* operon), cytochrome oxidase (the *cyo* operon), and several genes of menaquinone (the *menFDHBCE* operon), ubiquinone, and heme synthesis. The *speB* mutant also had fewer transcripts for all genes of iron acquisition, which is consistent with diminished oxidative energy generation. A *speB* mutant had fewer transcripts for *ptsH* (Hpr of the phosphotransferase system of carbohydrate transport), *ptsG* (enzyme IIBC component of glucose transport), *gltA* (citrate synthase), *sdhA* (a succinate dehydrogenase subunit), and *sucC* (a succinyl-CoA synthetase subunit), and their loss has been shown to either impair or eliminate W3110 surface motility (10). The cumulative effect of these differences could account for the motility defect in the *speB* mutant.

## Discussion

Several aspects of polyamine metabolism affect PDSM in *E. coli*. Strains with loss of one or two polyamine synthesis genes ― *speA* and *speB* single mutants and a *speC speF* double mutant ― resulted in PDSM defects. This observation is remarkable because nine enzymes and several pathways are involved in polyamine synthesis. Our putrescine synthesis mutants had no growth defect which is not unexpected since deletions of eight biosynthetic genes were required to show a strong phenotype: impaired aerobic growth and elimination of anaerobic growth (3). Strains with defects in the two most active putrescine transport systems had altered PDSM. The specific defects suggest the involvement of extracellular putrescine and arginine in intracellular putrescine homeostasis. The PDSM defect of putrescine transport mutants not only implicates extracellular putrescine but also suggests secretion of putrescine which is not added to the medium. The PDSM defect of *speA* and *speB* mutants implicates extracellular arginine because SpeA is mostly periplasmic (27). Defective putrescine catabolism also affected PDSM. The role of both putrescine synthesis and degradation suggests rapid adjustments to fluctuating putrescine concentrations and environmental conditions: putrescine catabolism is a core response to several stresses (19).

Results from an RNAseq analysis showed that transcription of the *fim* operon was reduced in the *speB* mutant. Of the known transcription factors that control the *fim* operon, only the gene for H-NS had fewer transcripts. In contrast, transcripts for other relevant transcription factor genes were elevated in the *speB* mutant. Our results are consistent with H-NS mediating putrescine-dependent control, but we cannot exclude a general regulatory dysfunction mediated by altered nucleoprotein complexes at the *fim* operon promoter. The elevated transcripts for some transcription factors may be inhibitory.

The transcriptomic results showed that the *speB* mutant had more transcripts for genes whose products will maintain intracellular putrescine, which suggests compensatory mechanisms for low intracellular putrescine (Figure 11). The *speB* mutant also had more transcripts for magnesium and phosphate transport genes. The elevated transcripts for magnesium transport genes could compensate for low putrescine to the extent that magnesium can directly replace putrescine, e.g., binding to nucleotides and RNA (28). Two recent studies observe an inverse correlation between intracellular magnesium and polyamines in *Salmonella* and proposed that either magnesium or the polyamines maintained an overall divalent cation homeostasis (29, 30). The rationale for an increase in anionic phosphate assimilation is less clear, although the major phosphate permease transports metal-phosphates, including magnesium-phosphate (31), which could help to compensate for low putrescine. Regardless of the rationale, the substantial changes in gene expression due to low putrescine suggest multiple compensatory mechanisms.

A major observation of the RNAseq analysis is that loss of *speB* lowered transcripts for many genes of energy metabolism and all genes of iron acquisition. This gene expression pattern suggests that low putrescine diverts metabolism away from energy generation and toward putrescine synthesis, i.e., a prioritization of putrescine synthesis over aerobic energy metabolism. Oxidative energy metabolism and putrescine synthesis compete for α-ketoglutarate and the metabolic diversion could also be a potential compensatory mechanism for low putrescine. Another way of looking at this relationship is that intracellular putrescine positively correlates with oxidative energy metabolism. Deletion of *ptsH*, *ptsG*, *gltA*, *sdhA*, and *sucC* have been shown to impair W3110 PDSM which suggests that a major, possibly the primary, effect of low putrescine is reduced energy generation.

In contrast to our results, a thorough and well-designed study showed that *E. coli* PDSM requires spermidine instead of putrescine (8). The differences between the studies were strain backgrounds, media (glucose-LB vs glucose-tryptone), and incubation temperature (33⁰ vs 37⁰). We found PDSM results for 37⁰ incubations were highly variable and resulted in numerous fast-moving variants. Since growth at the higher temperature is faster, and faster growth is associated with higher putrescine (32), we propose that growth at 37⁰ results in an inhibitory putrescine concentration. In this case, spermidine would be stimulatory because of its known inhibition of the putrescine biosynthetic enzymes SpeA and SpeC (33, 34) and subsequent reduction of intracellular putrescine to an optimal concentration. Regardless of the explanation, strains differences are not unexpected because of decades of strain passage for the strains employed with resulting laboratory evolution that can affect any of the numerous factors that control the intracellular putrescine concentration.

The transcript changes observed in our study were different from those in the only other systematic study of putrescine’s effect on *E. coli* gene expression (6). Both studies had almost identical experimental designs: 1.0- or 1.14-mM putrescine was added to a *speB* or *speB speC* mutant, respectively. Differences in strains, growth media (minimal media vs tryptone-containing media), and growth temperature (33⁰ vs 37⁰) could account for the discrepancy in results. The most crucial difference could be the growth media: Igarashi and colleagues grew bacteria in a minimal medium, while we grew bacteria in an amino acid-containing medium that was required for surface motility. We propose that the slower growth in minimal medium, possibly correlated with higher guanosine tetraphosphate which is associated with slower growth, counters and obscures many growth-stimulatory putrescine effects. In contrast, the growth stimulation from the amino acids in our medium would not counter putrescine effects and would essentially allow detection of effects caused solely by changes in putrescine concentrations. Regardless of the explanation, common results can be interpreted as core responses to putrescine limitation. Both studies observed that low putrescine increased transcripts for genes of arginine synthesis and magnesium transport, and reduced transcripts for the *sdhCDAB*-*sucABCD* operon which codes for three TCA cycle enzymes. The link between putrescine and arginine synthesis is probably mediated by the ArgR repressor because *argR* transcripts are 2.5-fold lower (FDR = 0.0004) in the *speB* mutant (Supplemental Table 1).

Several results implicate putrescine in *E. coli* virulence during urinary tract infections. First, pili-dependent binding to the urothelium is an essential step for uropathogenic *E. coli* (35). Second, putrescine is present in urine from infected, but not healthy, individuals (36, 37). Third, urinary putrescine can result from uropathogenic *E. coli* growth in urine (37) and urothelial pathophysiology (38). Putrescine’s possible role in virulence is seemingly contradicted by the observation that a *speB* mutant outcompetes the parental CFT073 in the bladder of mice, i.e., less putrescine synthesis enhanced virulence (39). A majority of uropathogenic *E. coli* are associated with phylogenetic group B2―including CFT073, whereas most *E. coli* lab strains, such as W3110, are group A. Group B2 strains lack genes for the major putrescine catabolic pathway and have significantly higher transcripts for genes coding for SpeA and SpeB (18). Both observations suggest that the cytoplasmic putrescine concentration is higher in group B2 strains. We propose that the putrescine concentration in group B2 strains is normally in the inhibitory range for pili synthesis, and that loss of SpeB reduces putrescine to a level that stimulates pili synthesis. Regardless of the explanation, putrescine control of pili synthesis and other processes in uropathogenic *E. coli* are worth examination.

## Experimental Procedures

### Strains

All strains used for growth rate determinations and motility assays were derivatives of *E. coli* K-12 strain W3110 and are listed in Table 3. Great variations exist in standard lab strains, including W3110, from different labs. We initially tested nine strains derived from *E. coli* K-12, mostly from the Coli Genetic Stock Center (9), and chose our lab strain because of relative genetic stability and more quantitatively reproducible results. Our strain originated from the lab of Jon Beckwith in the 1960s via the lab of Boris Magasanik where it had been stored in a room temperature stab until the mid-1980s, at which time it was revived and frozen at −80⁰ C. To construct the mutant strains, the altered allele was obtained from the Keio collection in which the gene of interest had been deleted and replaced with an antibiotic resistance gene (40). The marked deletion allele was transferred into W3110 by P1 transduction (41). The antibiotic gene was removed as described which generated an in-frame deletion (42).

**Table 3.**
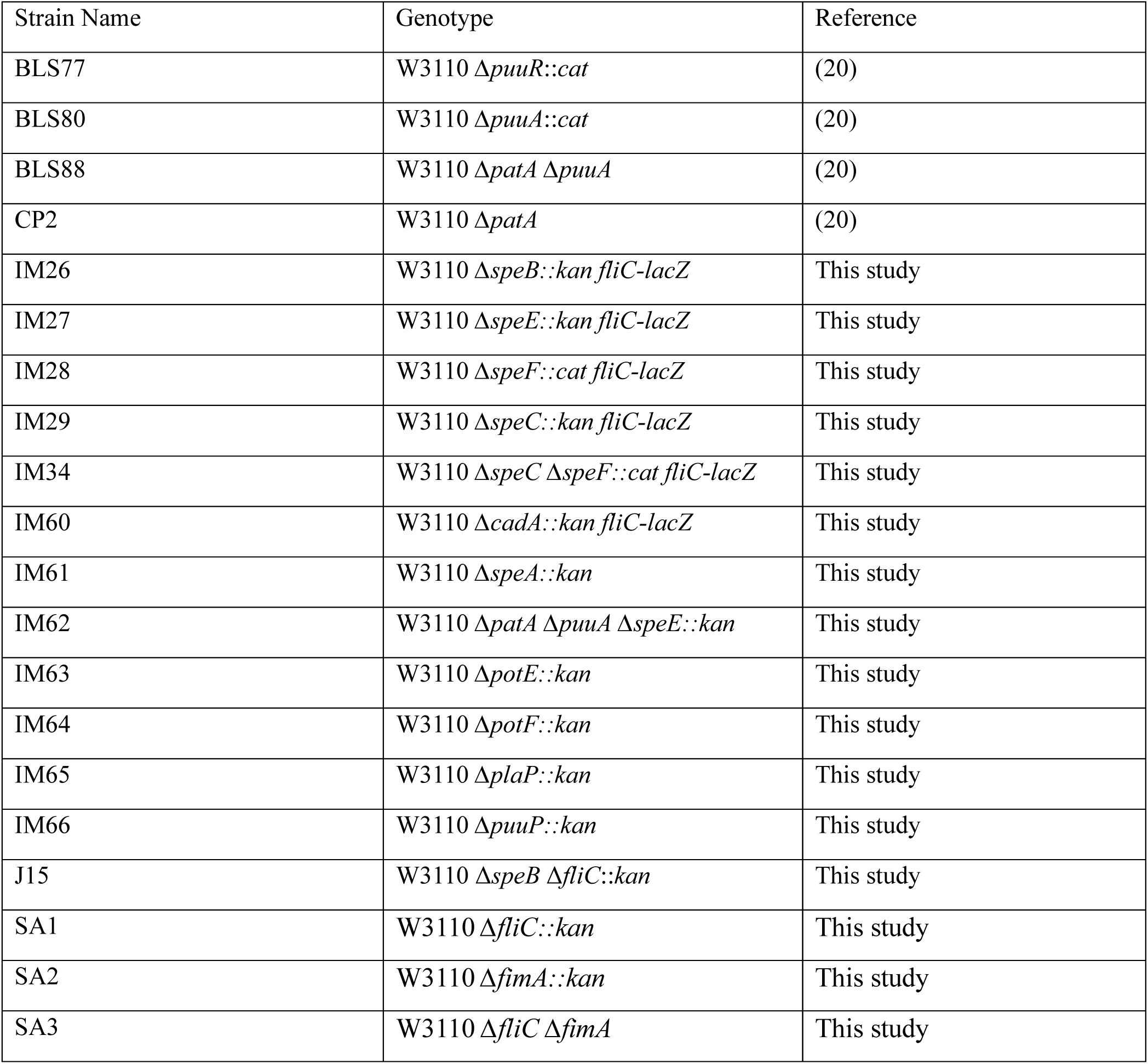
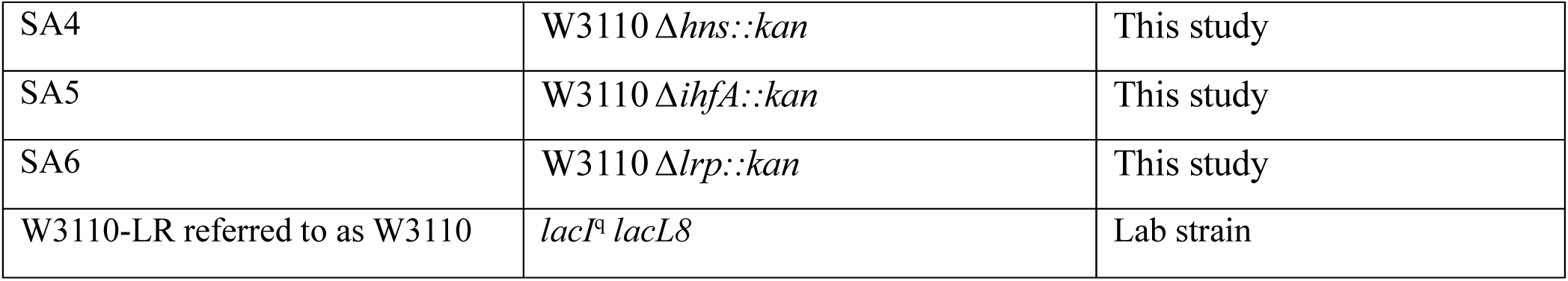
Strains.

### Media and Growth Conditions

For P1 transductions and plasmid transformation experiments, cells were grown in standard LB liquid medium (1% tryptone, 1% NaCl and 0.5% yeast extract, pH 7.0) at 37**°**C. Antibiotics were used for selection at concentrations of 25 µg/mL (both chloramphenicol and kanamycin). For growth analysis, bacteria were grown in liquid motility medium (0.5% glucose, 1% tryptone, and 0.25% NaCl) which we refer to as GT medium. Starter cultures (typically 6–12 h incubation) were grown in GT medium, harvested by centrifugation, washed twice with phosphate buffered saline and re-suspended in GT medium before inoculation. The cells were then grown at 37° C in aerobic conditions (shaking at 240 rpm) and the turbidity was measured every thirty minutes. Cell growth was measured in Klett units using a Scienceware Klett colorimeter with a KS-54 filter. 100 Klett units represent an OD_600_ value of about 0.7.

### Surface motility assay

For a standard surface motility assay, single colonies from fresh plates (streaked out from frozen stocks a day before) were inoculated in GT medium for six hours at 37° C in aerobic conditions (shaking at 240 rpm). 30 ml of autoclaved GT medium with 0.45% agar (Eiken, Tokyo, Japan) was poured into sterile polystyrene petri-dish (100 mm X 15 mm) and allowed to solidify at room temperature for approximately six hours. Then, 1 µL of the pre-motility growth medium was inoculated at the center of the agar plate and incubated at 33**°** C for 36 hrs. Each experiment was performed in triplicate and pictures were taken after 36 hours. Surface motility was extremely sensitive to humidity. Opening the incubator before 36 hours resulted in movement cessation. The surface motility assay was performed at 33⁰ C because results from incubations at 37⁰ C were highly variable and the cultures more frequently generated genetically stable fast-moving variants.

### Swim assay

The media and culturing for swimming motility is identical to that for surface motility, except that the plates contained 0.25% agar and were solidified at room temperature for about one hour before inoculation. Then, 1 µL of the culture was stabbed in the center of this swim agar plate and incubated at 33**°** C for 20 hours. Each experiment was performed in triplicate.

### Transmission electron microscopy

Cells from surface motility plates were collected and fixed with 2.5% glutaraldehyde. Bacteria were absorbed onto Foamvar carbon-coated copper grids for 1 min. Grids were washed with distilled water and stained with 1% phosphotungstic acid for 30 s. Washed and stained grids were dried at 37⁰C for 10 minutes. Samples were viewed on a JEOL 1200 EX transmission electron microscope at University of Texas Southwestern Medical Center (Figure 2) and University of Texas at Dallas (Figure 8).

### ELISA analysis

Cells were grown overnight in 5 mL GT media followed by a 2-hour growth in GT media with 0.0, 0.1, 1.0, or 4.0 mM putrescine. Cells were then lysed using a 24-gauge needle. The cell lysate was diluted to a concentration of 1 mg/mL protein (based on A280) in pH 9.6 coating buffer (3 g Na_2_CO_3_, 6 g NaHCO_3_, 1000 mL distilled water) and then coated onto the walls of a 96-well plate by incubating overnight at 4°C. The wells were rinsed 3 times with PBS and further blocked using coating buffer with 1% bovine albumin overnight at 4°C. The following day, wells were rinsed 3 times with PBS, and primary antibodies to FimA or RpoD were added following supplier directions. After a 2-hour room temperature incubation, secondary antibodies conjugated to HRP were added and incubated for a further two hours. HRP was activated using the TMB substrate kit (Thermo-34021) following the provided protocol. Expression was read on a BioTek plate reader based on the provided kit protocol. Data was blanked to wells only treated with bovine albumin then normalized to RpoD. Data was generated from technical and biological triplicates.

### DNA isolation and genome assembly and annotation for W3110

DNA was isolated based on previous established protocols (43). Short reads were sequenced at the SeqCenter (Pittsburgh, Pennsylvania) using Illumina tagmentation-based and PCR-based DNA prep and custom IDT indices targeting inserts of 280 bp without further fragmentation or size selection steps. The Illumina NovaSeq X Plus sequencer was ran producing 2X151 paired-end reads. Demultiplexing, quality control, and adapter trimmer was performed with bcl-convert (v4.2.4). Total Illumina Reads (R1+R2): 4059078 with 553087045 bps >Q30.

Long reads were prepared using the PCR-free Oxford Nanopore Technologies ligation sequencing kit with the NEBNext companion module. No fragmentation or size selection was performed. Long read libraries were performed using R10.4.1 flow cell. The 400bps sequencing mode with a minimum read length of 200bp were selected. Gupply (v6.5.7) was used at the super-accurate basecalling, demultiplexing, and adapter removal. Total long reads: 53773 with 89.607 % of bps > Q20.

To generate the completed genome, porechop (v0.2.4) was used to trim residual adapter sequences from long reads. Flye (v2.9.2) was used to generate the *de novo* assembly under the nano-hq model. Reads longer than the estimated N_50_ based on a genome size of 6Mbp initiated the assembly. Subsequent polishing using the short read data was performed using Pilon (v1.24) under default parameters. Long read contigs with an average short read coverage of 15X or less were removed from the assembly. The assembled contig was confirmed to be circular via circulator (v1.5.5). Annotation was performed using prokka (v 1.14.6). Finally, statistics were recorded using QUAST (v5.2.0). The final genome contained 1 contig of 4750347 bp with a sequencing depth of 123.92x. The N_50_ was 4750347. This genome can be accessed via the accession number CP165600 or via the BioProject PRJNA1142534 via NCBI. The completed genome has an estimated average nucleotide identity of 99.9694 to the W3110 genome deposited to NCBI (genome assembly ASM1024v1).

### RNA isolation and quality control

Cells were grown for 2 hours in 1 mL GT media. 60 µL were then added to 1 mL of fresh GT, with or without 1 mM putrescine, and grown for another two hours. After growth, the cells were centrifuged, and frozen at −80⁰C. The isolation and analysis protocol has been previously published (18). Cell pellets were thawed, resuspended in 0.7 mL of buffer RLT (Qiagen RNeasy kit), and mechanically lysed using a bead beater (FastPrep-24 Classic from MP Biomedical) set at the highest setting for three 45-s cycles with a 5-min rest period on ice between cycles. Cell lysates were used to isolate RNA using a Qiagen RNeasy mini kit. RNA was quantified, DNase treated, and re-quantified on a nanodrop. RNA that passed the first quality check was analyzed on RNA-IQ (Qubit). RNA with an IQ higher than 7.5 was used. RNA that passed both checks were run on an agarose gel to check for RNA integrity. RT-qPCR was used to ensure there was no DNA contamination. RNA libraries that passed all these quality checks were submitted to the genomics core facility at The University of Texas at Dallas for RNA sequencing. The core performed rRNA removal (RiboMinus Transcriptome Isolation Kit or RiboCop bacterial rRNA depletion—Lexogen), library preparation (Stranded Total RNA Prep—Illumina), and single-end Illumina sequencing.

### Reverse transcription quantitative PCR

RNA was isolated as described above. One microgram of RNA was reverse transcribed using LunaScript RT Super Mix (NEB E3010) and another microgram was subjected to the same reaction as the reverse transcribed RNA, except without reverse transcriptase (negative control) following manufacturer instructions. The cDNA was then diluted to 10 ng/µL for qPCR. PCR reactions were composed of 10 ng cDNA, 8 µL of nuclease free water, 10 nM primers (rpoD: Forward: TCGTGTTGAAGCAGAAGAAGCG; Reverse: TCGTCATCGCCATCTTCTTCG) (fimA: Forward: ATGGTGGGACCGTTCACTTT; Reverse: GGCAACAGCGGCTTTAGATG), and PowerUp SYBR Green 2X master mix for qPCR (ThermoFisher A25777) in a 20 µL reaction. A Quantstudio 3 qPCR machine was used to generate critical threshold (Ct) values. The Ct values were then analyzed using the 2^-ΔΔC^_T_ method (44) using *rpoD* as a library control.

### RNAseq Analysis

Transcripts were aligned to the W3110 genome and CDS (CP165600) available on NCBI using CLC genomics workbench (Qiagen version 22.00) for RNA-seq analysis. The gene expression counts generated as this output were used to perform differentially expressed gene (DEG) analysis using EdgeR (version 3.38.4) to analyze read counts (45, 46). DEGs had an adjusted P-value less than 0.05.

## Data availability

The raw RNA seq reads were deposited in GenBank: BioProject accession number PRJNA1126736. The genome was deposited in GenBank: BioProject accession number PRJNA1142534. The genome accession number is CP165600.

## Acknowledgements

This work was funded through private donations. PEZ received endowment support from the Cain Foundation through the Felecia and John Cain Distinguished Chair in Women’s Health in honor of Philippe E. Zimmern, MD. The authors declare no potential conflicts of interests.

The authors (JH, SH, GV, and LR) would like to thank Prof. Kelli Palmer of The University of Texas at Dallas for the use of her space, supplies, and tissue homogenizer. JH would like to further thank Prof. Palmer for the use of her CLC genomics program. The authors would like to thank the Genome Center at The University of Texas at Dallas for the services to support our research. The authors would like to thank the Olympus Discovery Center/Imaging Core facility at UT Dallas for providing equipment and support (Figure 8), and the UT Southwestern Medical Center for electron microscopy (Figure 2).

---

Supplemental Figure 1: Growth and swimming of polyamine anabolic mutants.

Supplemental Table 1. Differential gene expression analysis of W3110 and Δ*speB*. J15 refers to the Δ*speB* mutant. In the tab labeled “W3110_0_vs_J15_1_ Putrescine” W3110 was grown without putrescine, J15 was grown with 1.0 mM putrescine; positive fold change values refer to increases in J15 gene expression.

Supplementary Figure 2. Expression graphs comparing the logCPM of the transcriptomes of the average of the three replicates of W3110 with and without 1 mM putrescine and the *speB* mutant with and without 1 mM putrescine. A regression line was calculated and the correlation between each set of transcriptomes noted on the graph. Higher *R*^2^ values indicate greater similarity between the transcriptomes.

Supplemental Figure 3. Phase variation in parental W3110, W3110 Δ*speB*, W3110 Δ*hns*, and MG1655

## References

1. Tabor CW, Tabor H. 1984. Polyamines. Annual review of biochemistry 53:749–90.

2. Tabor CW, Tabor H. 1984. Polyamines. Annu Rev Biochem 53:749–90.

3. Chattopadhyay MK, Tabor CW, Tabor H. 2009. Polyamines are not required for aerobic growth of *Escherichia coli*: preparation of a strain with deletions in all of the genes for polyamine biosynthesis. J Bacteriol 191:5549–52.

4. Igarashi K, Kashiwagi K. 2000. Polyamines: mysterious modulators of cellular functions. Biochem Biophys Res Commun 271:559–64.

5. Igarashi K, Kashiwagi K. 2015. Modulation of protein synthesis by polyamines. IUBMB Life 67:160–9.

6. Yoshida M, Kashiwagi K, Shigemasa A, Taniguchi S, Yamamoto K, Makinoshima H, Ishihama A, Igarashi K. 2004. A unifying model for the role of polyamines in bacterial cell growth, the polyamine modulon. J Biol Chem 279:46008–13.

7. Armbruster CE, Hodges SA, Mobley HL. 2013. Initiation of swarming motility by *Proteus mirabilis* occurs in response to specific cues present in urine and requires excess L-glutamine. J Bacteriol 195:1305–19.

8. Kurihara S, Suzuki H, Tsuboi Y, Benno Y. 2009. Dependence of swarming in *Escherichia coli* K-12 on spermidine and the spermidine importer. FEMS Microbiol Lett 294:97–101.

9. Ambagaspitiye S, Sudarshan S, Hogins J, McDill P, De Nisco NJ, Zimmern PE, Reitzer L. 2019. Fimbriae and flagella mediated surface motility and the effect of glucose on nonpathogenic and uropathogenic *Escherichia coli* BioRxiv doi.org:10.1101/840991

10. Sudarshan S, Hogins J, Ambagaspitiye S, Zimmern P, Reitzer L. 2021. The nutrient and energy pathway requirements for surface motility of nonpathogenic and uropathogenic *Escherichia coli*. J Bacteriol. doi:10.1128/jb.00467-20.

11. Yokota T, Gots JS. 1970. Requirement of adenosine 3’, 5’-cyclic phosphate for flagella formation in *Escherichia* coli and *Salmonella typhimurium*. J Bacteriol 103:513–6.

12. Conway C, Beckett MC, Dorman CJ. 2023. The DNA relaxation-dependent OFF-to-ON biasing of the type 1 fimbrial genetic switch requires the Fis nucleoid-associated protein. Microbiology (Reading). 169(1):doi:10.1099/mic.0.001283.

13. Kelly A, Conway C, T OC, Smith SG, Dorman CJ. 2006. DNA supercoiling and the Lrp protein determine the directionality of fim switch DNA inversion in *Escherichia coli* K-12. J Bacteriol 188:5356–63.

14. O’Gara JP, Dorman CJ. 2000. Effects of local transcription and H-NS on inversion of the fim switch of *Escherichia coli*. Mol Microbiol 36:457–66.

15. Tabor CW, Tabor H. 1985. Polyamines in microorganisms. Microbiol Rev 49:81–99.

16. Igarashi K, Ito K, Kashiwagi K. 2001. Polyamine uptake systems in *Escherichia coli*. Res Microbiol 152:271–8.

17. Kurihara S, Suzuki H, Oshida M, Benno Y. 2011. A novel putrescine importer required for type 1 pili-driven surface motility induced by extracellular putrescine in *Escherichia coli* K-12. J Biol Chem 286:10185–92.

18. Hogins J, Xuan Z, Zimmern PE, Reitzer L. 2023. The distinct transcriptome of virulence-associated phylogenetic group B2 *Escherichia coli*. Microbiol Spectr. doi:10.1128/spectrum.02085-23e0208523. doi:10.1128/spectrum.02085-23.

19. Schneider BL, Hernandez VJ, Reitzer L. 2013. Putrescine catabolism is a metabolic response to several stresses in *Escherichia coli*. Mol Microbiol 88:537–50.

20. Schneider BL, Reitzer L. 2012. Pathway and enzyme redundancy in putrescine catabolism in *Escherichia coli*. J Bacteriol 194:4080–8.

21. Fukuchi J, Kashiwagi K, Yamagishi M, Ishihama A, Igarashi K. 1995. Decrease in cell viability due to the accumulation of spermidine in spermidine acetyltransferase-deficient mutant of *Escherichia coli*. J Biol Chem 270:18831–5.

22. Sakamoto A, Sahara J, Kawai G, Yamamoto K, Ishihama A, Uemura T, Igarashi K, Kashiwagi K, Terui Y. 2020. Cytotoxic mechanism of excess polyamines functions through translational repression of specific proteins encoded by polyamine modulon. Int J Mol Sci. 21(7):doi:10.3390/ijms21072406.

23. Schwan WR. 2011. Regulation of *fim* genes in uropathogenic *Escherichia coli*. World J Clin Infect Dis 1:17–25.

24. Karp PD, Ong WK, Paley S, Billington R, Caspi R, Fulcher C, Kothari A, Krummenacker M, Latendresse M, Midford PE, Subhraveti P, Gama-Castro S, Muniz-Rascado L, Bonavides-Martinez C, Santos-Zavaleta A, Mackie A, Collado-Vides J, Keseler IM, Paulsen I. 2018. The EcoCyc Database. EcoSal Plus. 8(1):doi:10.1128/ecosalplus.ESP-0006-2018.

25. Sugiyama Y, Nakamura A, Matsumoto M, Kanbe A, Sakanaka M, Higashi K, Igarashi K, Katayama T, Suzuki H, Kurihara S. 2016. A novel putrescine exporter SapBCDF of *Escherichia coli*. J Biol Chem 291:26343–26351.

26. Keseler IM, Mackie A, Santos-Zavaleta A, Billington R, Bonavides-Martínez C, Caspi R, Fulcher C, Gama-Castro S, Kothari A, Krummenacker M, Latendresse M, Muñiz-Rascado L, Ong Q, Paley S, Peralta-Gil M, Subhraveti P, Velázquez-Ramírez DA, Weaver D, Collado-Vides J, Paulsen I, Karp PD. 2016. The EcoCyc database: reflecting new knowledge about *Escherichia coli* K-12. Nucleic Acids Research 45:D543–D550.

27. Tabor H, Tabor CW. 1969. Partial separation of two pools of arginine in *Escherichia coli*; preferential use of exogenous rather than endogenous arginine for the biosynthesis of 1,4-diaminobutane. J Biol Chem 244:6383–7.

28. Miyamoto S, Kashiwagi K, Ito K, Watanabe S, Igarashi K. 1993. Estimation of polyamine distribution and polyamine stimulation of protein synthesis in *Escherichia coli*. Arch Biochem Biophys 300:63–8.

29. Iwadate Y, Golubeva YA, Slauch JM. 2023. Cation homeostasis: Coordinate regulation of polyamine and magnesium levels in *Salmonella*. mBio 14:e0269822.

30. Duprey A, Groisman EA. 2020. DNA supercoiling differences in bacteria result from disparate DNA gyrase activation by polyamines. PLoS Genet 16:e1009085.

31. van Veen HW, Abee T, Kortstee GJ, Konings WN, Zehnder AJ. 1994. Translocation of metal phosphate via the phosphate inorganic transport system of *Escherichia coli*. Biochemistry 33:1766–70.

32. Tweeddale H, Notley-McRobb L, Ferenci T. 1998. Effect of slow growth on metabolism of *Escherichia coli*, as revealed by global metabolite pool (“metabolome”) analysis. J Bacteriol 180:5109–16.

33. Applebaum DM, Dunlap JC, Morris DR. 1977. Comparison of the biosynthetic and biodegradative ornithine decarboxylases of *Escherichia coli*. Biochemistry 16:1580–4.

34. Wu WH, Morris DR. 1973. Biosynthetic arginine decarboxylase from *Escherichia coli*. Purification and properties. J Biol Chem 248:1687–95.

35. Flores-Mireles AL, Walker JN, Caparon M, Hultgren SJ. 2015. Urinary tract infections: epidemiology, mechanisms of infection and treatment options. Nat Rev Microbiol 13:269–84.

36. Puebla-Barragan S, Renaud J, Sumarah M, Reid G. 2020. Malodorous biogenic amines in *Escherichia coli*-caused urinary tract infections in women-a metabolomics approach. Sci Rep 10:9703.

37. Satink HP, Hessels J, Kingma AW, van den Berg GA, Muskiet FA, Halie MR. 1989. Microbial influences on urinary polyamine excretion. Clin Chim Acta 179:305–14.

38. Keay SK, Birder LA, Chai TC. 2014. Evidence for bladder urothelial pathophysiology in functional bladder disorders. Biomed Res Int 2014:865463.

39. Alteri CJ, Smith SN, Mobley HL. 2009. Fitness of *Escherichia coli* during urinary tract infection requires gluconeogenesis and the TCA cycle. PLoS Pathog 5:e1000448.

40. Baba T, Ara T, Hasegawa M, Takai Y, Okumura Y, Baba M, Datsenko KA, Tomita M, Wanner BL, Mori H. 2006. Construction of *Escherichia coli* K-12 in-frame, single-gene knockout mutants: the Keio collection. Mol Syst Biol 2:2006 0008.

41. Miller JH. 1972. Experiments in Molecular Genetics. Cold Spring Harbor Laboratory, Cold Spring Harbor, NY.

42. Datsenko KA, Wanner BL. 2000. One-step inactivation of chromosomal genes in *Escherichia coli* K-12 using PCR products. Proc Natl Acad Sci U S A 97:6640–5.

43. Hogins J, Zimmern PE, Reitzer L. 2023. Genome sequences of seven clade B2 *Escherichia coli* strains associated with recurrent urinary tract infections in postmenopausal women. Microbiol Resour Announc doi:10.1128/mra.00035-23:e0003523.

44. Livak KJ, Schmittgen TD. 2001. Analysis of relative gene expression data using real-time quantitative PCR and the 2(-Delta Delta C(T)) Method. Methods 25:402–8.

45. McCarthy DJ, Chen Y, Smyth GK. 2012. Differential expression analysis of multifactor RNA-Seq experiments with respect to biological variation. Nucleic Acids Res 40:4288–97.

46. Robinson MD, McCarthy DJ, Smyth GK. 2010. edgeR: a Bioconductor package for differential expression analysis of digital gene expression data. Bioinformatics 26:139–40.

