## Supplemental figure 1 for "Control of pili synthesis and putrescine homeostasis in *Escherichia coli*"

A. Growth rates are the same for parental, Δ*speA*, Δ*speB*, and the Δ*speC* Δ*speF* double mutant strains when grown in glucose tryptone medium (liquid motility medium).

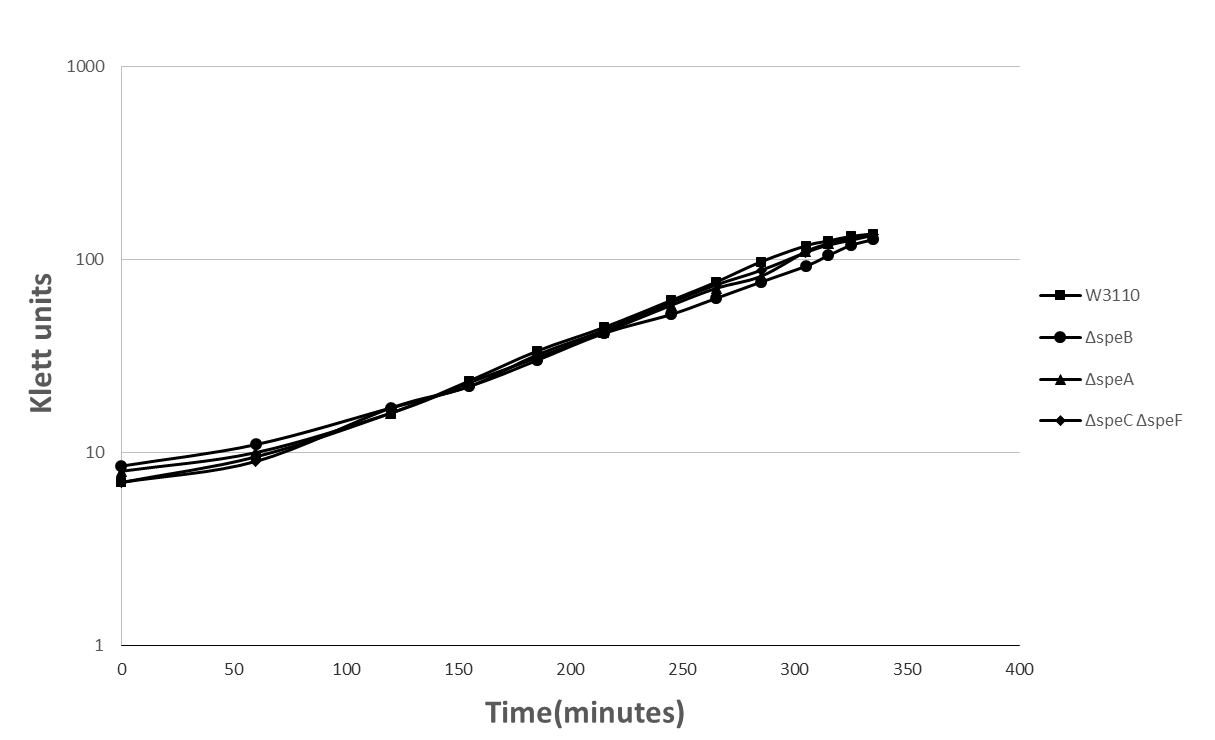

B. Loss of putrescine synthetic genes does not affect swimming motility.

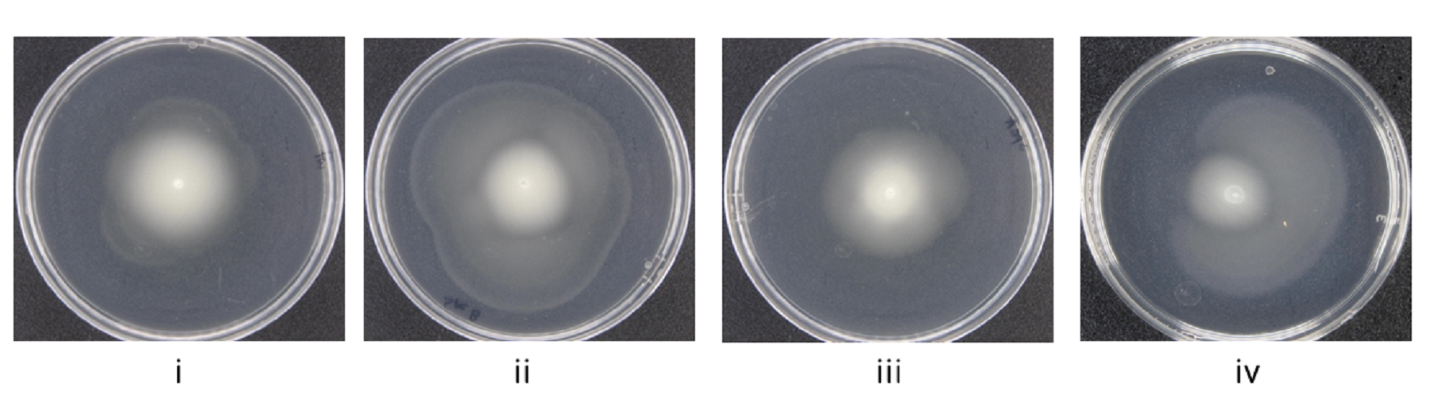

Parental W3110	Δ*speB*	Δ*speA*	Δ*speC* Δ*speF*
