## Supplemental figure 2 for "Control of pili synthesis and putrescine homeostasis in *Escherichia coli*"

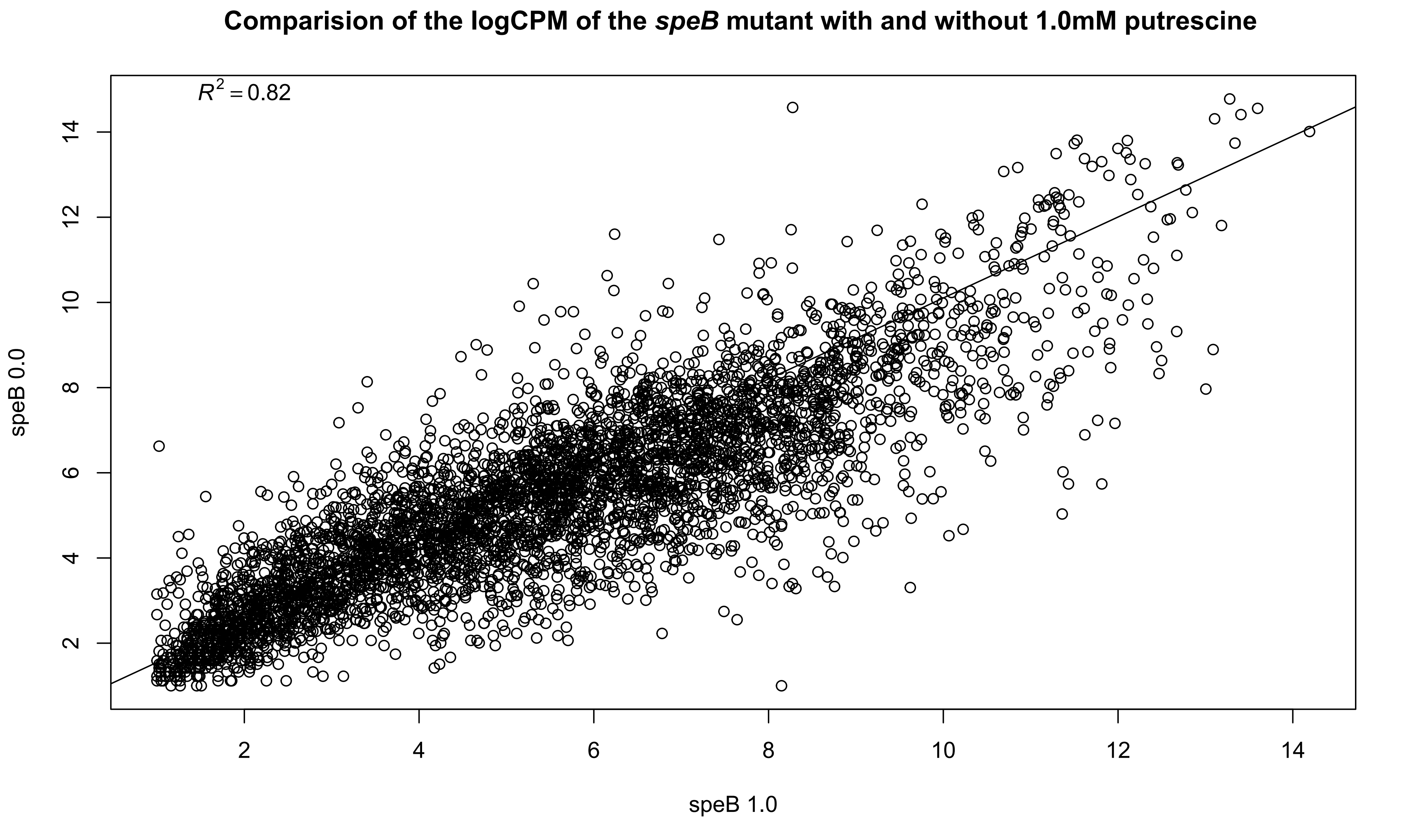

Comparision of the logCPM of the *speB* mutant with and without 1.0mM putrescine
$R^2 = 0.82$
speB 0.0
speB 1.0
2
4
6
8
10
12
14

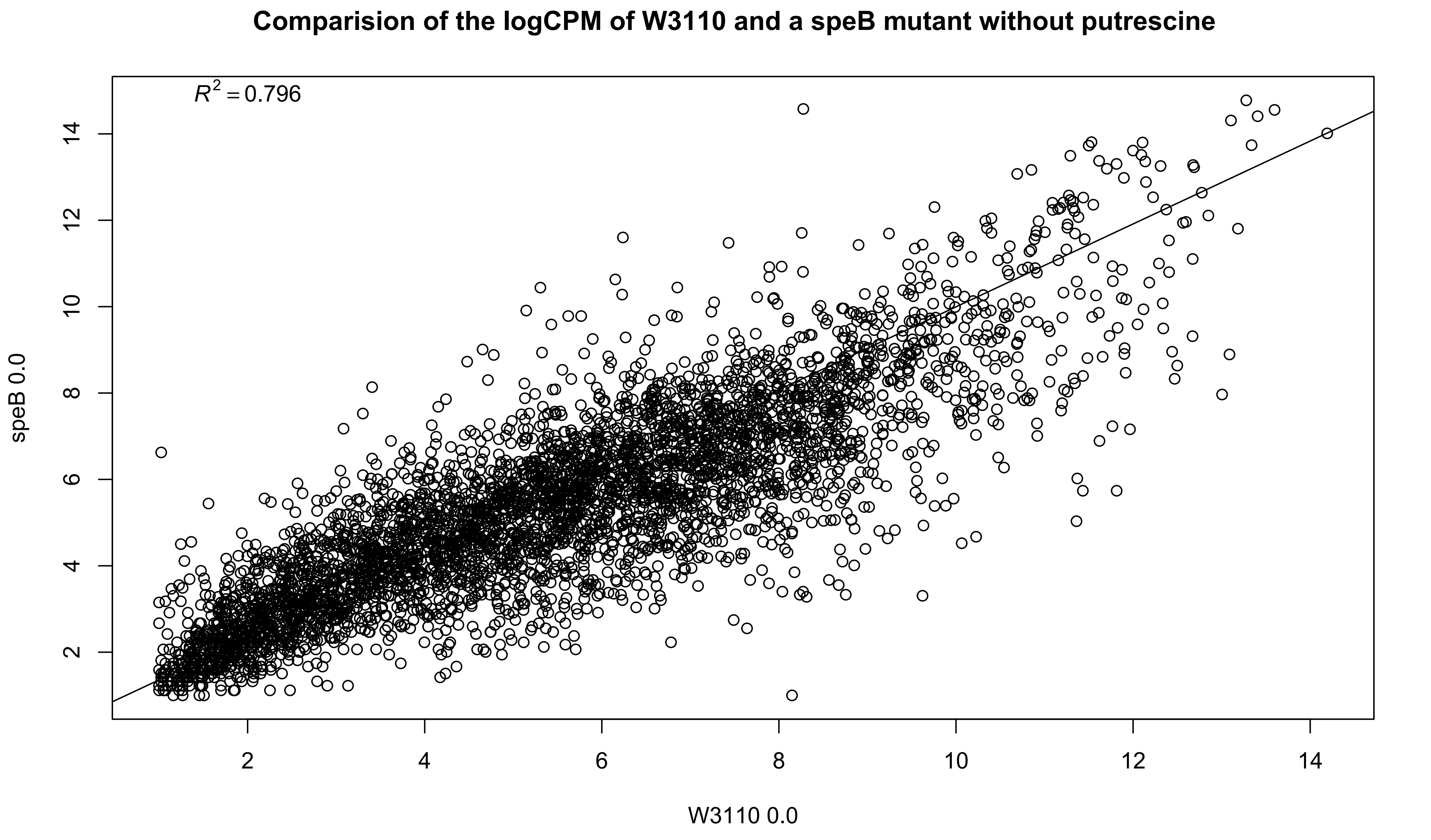
Comparision of the logCPM of W3110 and a speB mutant without putrescine
$R^2 = 0.796$
speB 0.0
W3110 0.0
2
4
6
8
10
12
14
2
4
6
8
10
12
14

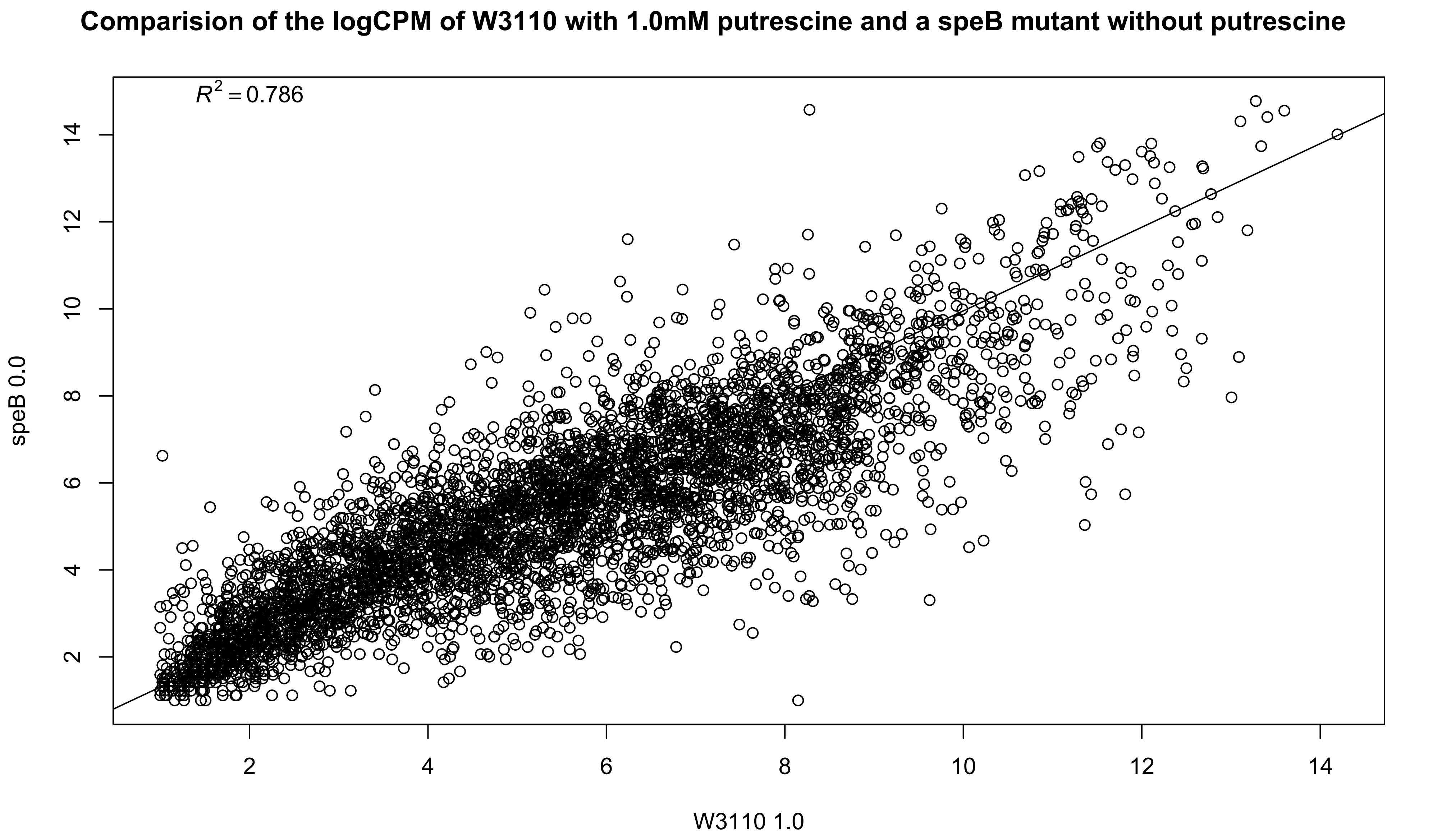

Comparision of the logCPM of W3110 with 1.0mM putrescine and a speB mutant without putrescine
$R^2 = 0.786$
speB 0.0
W3110 1.0
2
4
6
8
10
12
14

### Comparision of the logCPM of W3110 and a speB mutant with 1.0mM putrescine

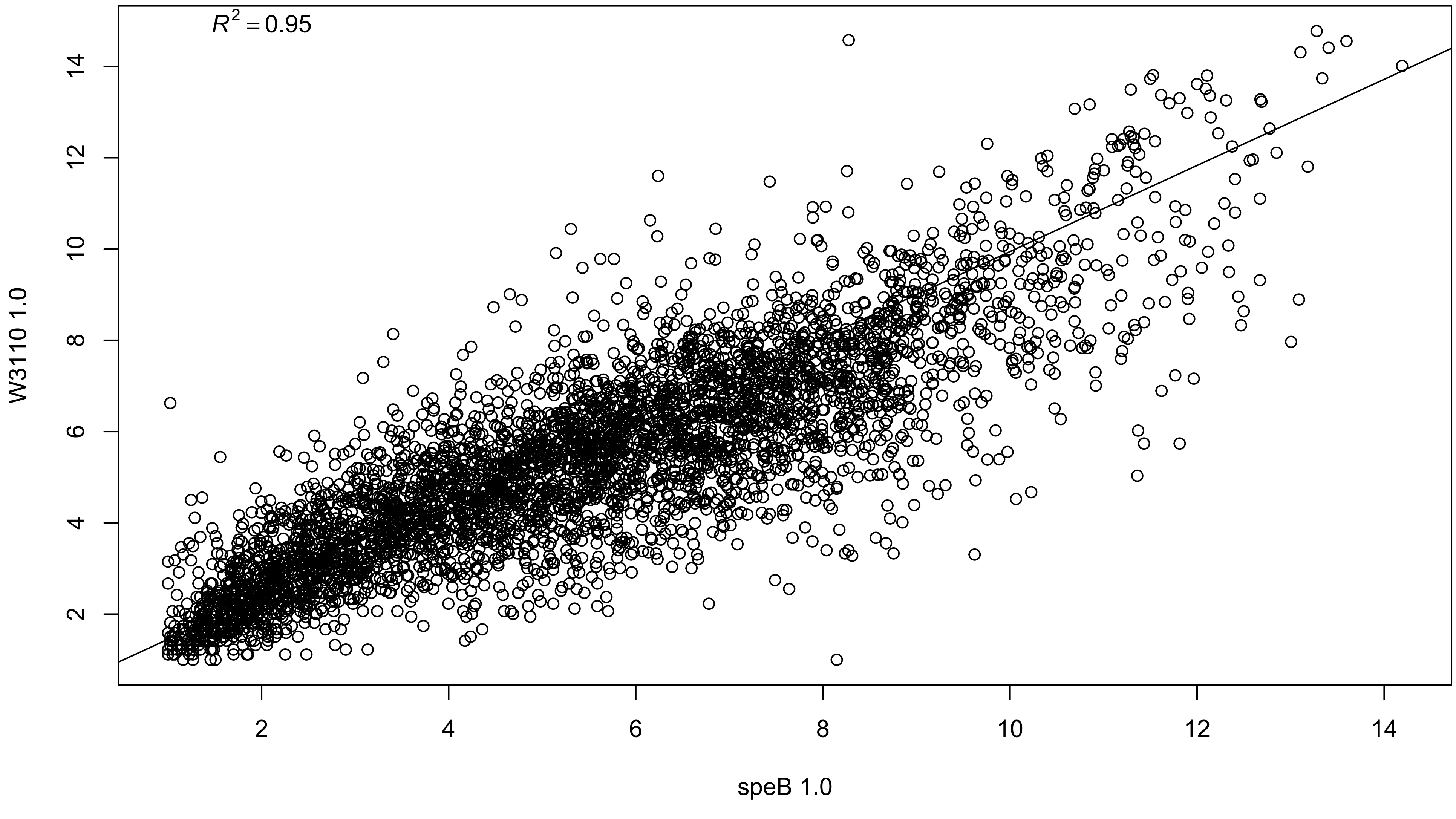

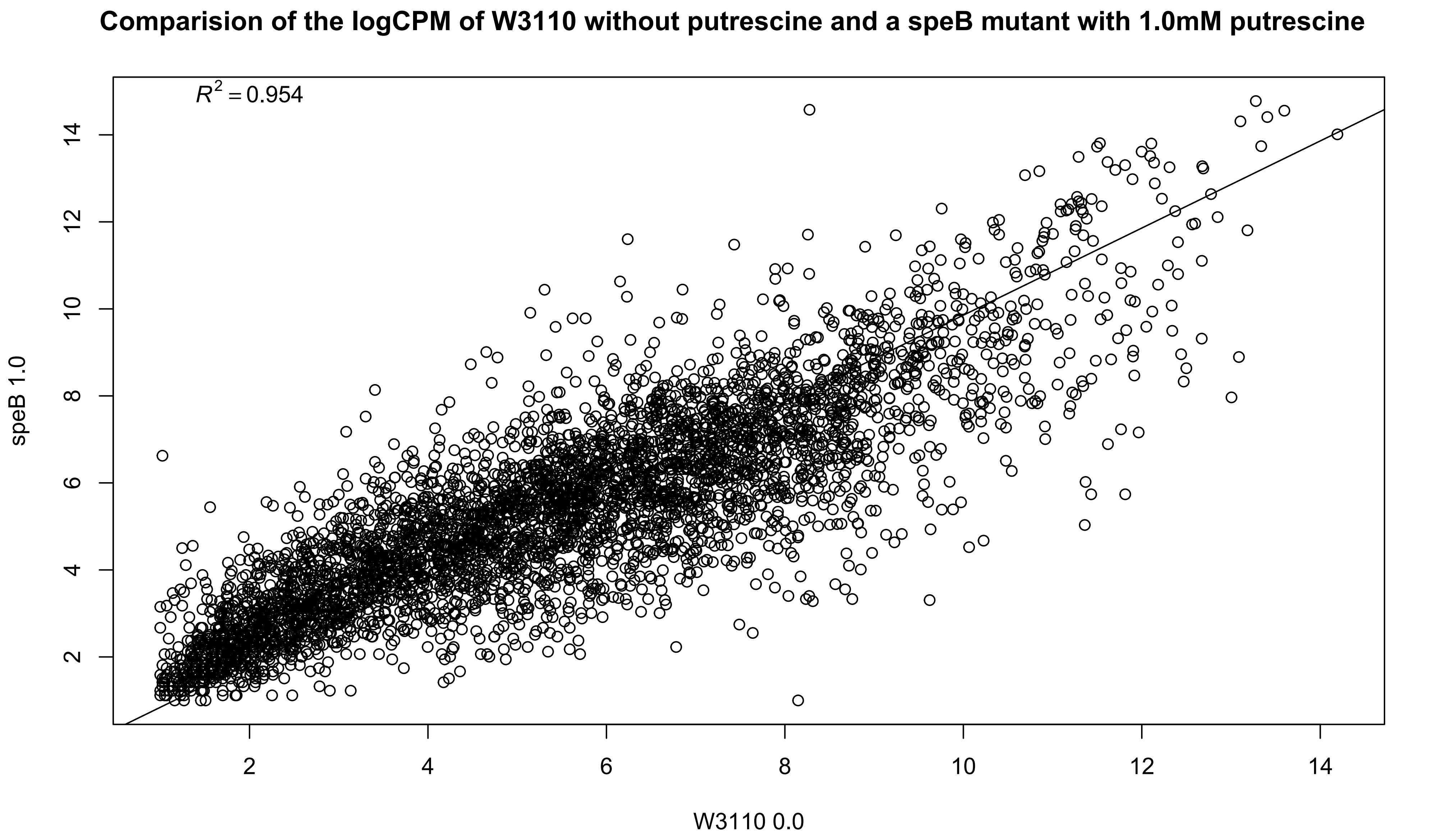

Comparision of the logCPM of W3110 without putrescine and a speB mutant with 1.0mM putrescine
$R^2 = 0.954$
speB 1.0
W3110 0.0
2
4
6
8
10
12
14

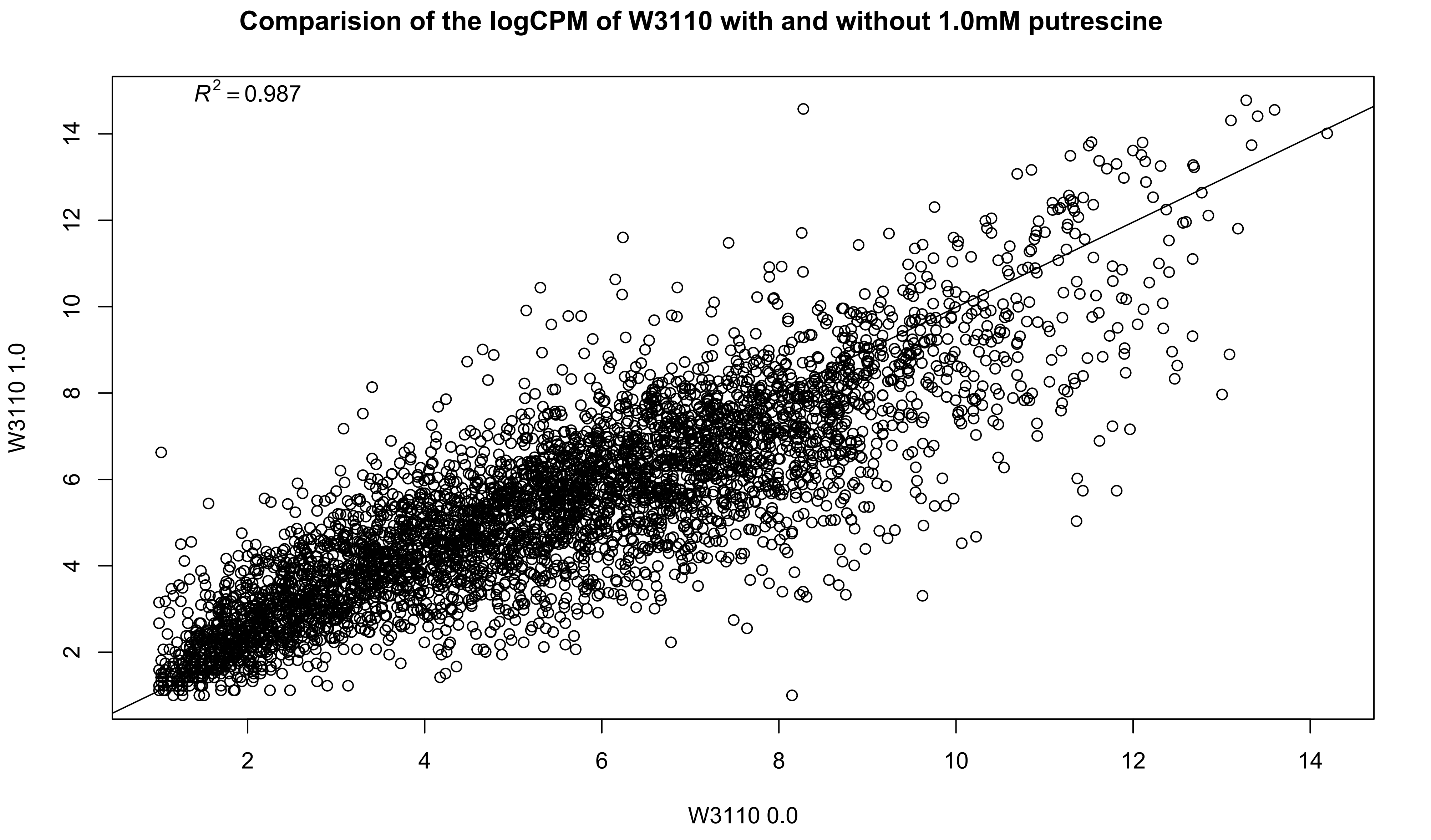
Comparision of the logCPM of W3110 with and without 1.0mM putrescine
$R^2 = 0.987$
W3110 1.0
W3110 0.0
2
4
6
8
10
12
14

Supplementary Figure 2 Legned

Supplementary Figure 2: Expression graphs comparing the logCPM of the transcriptomes of the average of the three replicates of the W3110 with and without 1 mM putrescine and the *speB* mutant with and without 1 mM putrescine. A regression line was calculated and the correlation between each set of transcriptomes noted on the graph. Higher $R^2$ values indicate greater similarity between the transcriptomes.
