## Supplemental Figure 3 for "Control of pili synthesis and putrescine homeostasis in *Escherichia coli*"

Supplemental Figure 3 Phase variation in parental W3110, W3110 Δ*speB*, W3110 Δ*hns* and MG1655

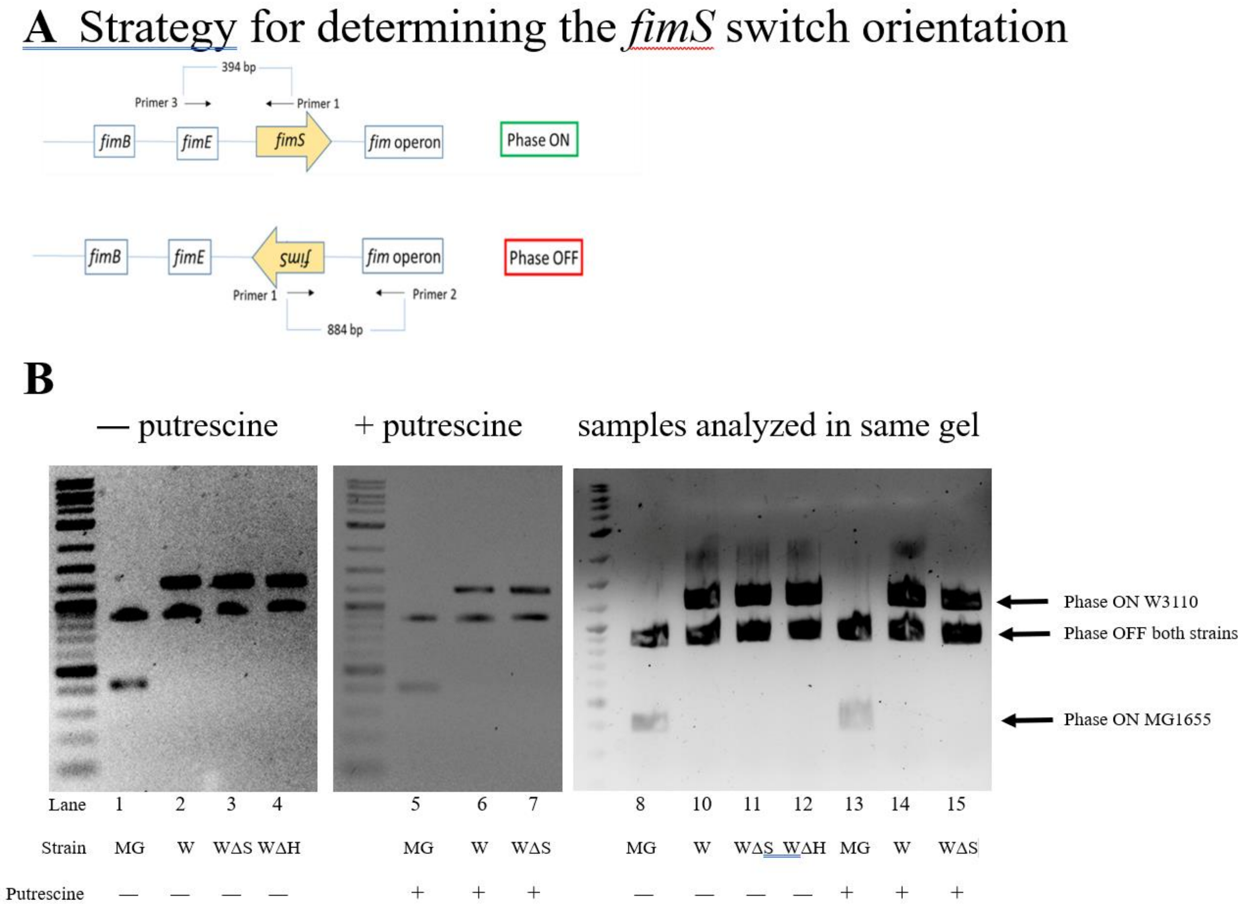


A. The diagram shows the genes for the FimB and FimE recombinases, the invertible *fimS* region which contains the promoter for the *fim* operon, and the *fim* operon which codes for the proteins of the type 1 pilus. Primer pairs 1-2 and 1-3 detect the *fimS* region in the phase OFF and ON orientations, respectively. The DNA sizes for phase OFF and ON are 884 and 394, respectively, for wild-type strains of *E. coli*, such as MG1655. Our lab strain of W3110 has an *IS1* element insertion in *fimE* which increases the size of the amplified DNA fragment. MG1655 was analyzed as a control. Primer 1 is 5'-CCGCGATGCTTTCCTCTATG-3'; primer 2 is 5'-TAATGACGCCCTGAAATTGC-3'; and primer 3 is 5'-TGCTAACTGGAAAGGCGCTG-3' (shown schematically).

B. Deletion of either *speB* or *hns* had no effect on *fimS* orientation. A possible explanation for loss of pili or PDSM in the *speB* or *hns* mutants is locking the *fimS* switch in phase OFF. However, loss of either *speB* or *hns* had no effect on *fimS* orientation in W3110 (lanes 2-4), and putrescine did not alter *fimS* orientation of the W3110 *ΔspeB* mutant (lanes 6 and 7). We conclude that loss of *speB* in W3110 did not phase-lock *fimS* in phase OFF. Also note that W3110, which has an insertion in *fimE*, is not locked in phase ON. The strains are MG1655 (MG), parental W3110 (W), W3110 *ΔspeB* (WΔS), and W3110 *Δhns* (WΔH).
